# Hologenomics of xylotrophic bivalves reveals a minimalist, remote-acting evolutionary strategy of wood digestion

**DOI:** 10.64898/2026.07.25.740686

**Authors:** Hao Song, Biyang Xu, Yang Guo, Cong Zhou, Meijie Yang, Yantao Liu, Zhaoshan Zhong, Chun-Yang Li, Xiaolin Tian, Yanyan Wang, Ron Flatau, Minxiao Wang, Tao Zhang, Daniel L. Distel, Yuanning Li

## Abstract

Wood constitutes the largest reservoir of biogenic carbon on Earth, yet remarkably few animals can exploit it. While terrestrial wood-feeders like termites rely on highly diverse gut microbiomes, xylotrophic marine bivalves have evolved a fundamentally different approach: a spatially segregated system where intracellular gill symbionts produce enzymes that act remotely within a nearly sterile cecum. However, the genetic and evolutionary basis of this unique symbiosis remains largely elusive. Here, we integrate hologenomics, transcriptomics, and biochemistry of a shallow-water shipworm (*Teredo navalis*) and a deep-sea borer (*Xyloredo* sp.). We find that despite diverging approximately 147 million years ago and occupying drastically different habitats, these bivalves maintain a strictly conserved ancestral karyotype and a shared genomic architecture for wood digestion. Our models reveal a clear host-symbiont division of labor. The host genome is specialized for lignin modification and targeted enzyme transport, whereas a highly streamlined symbiont community is responsible for core polysaccharide degradation. Central to this minimalist strategy is a lineage-specific GH5–GH6 dual-catalytic enzyme. By sharing amino acids across proximal binding pockets, this fusion protein unites endo- and exo-cellulase activities, enabling highly synergistic cellulose cleavage without the need for complex microbial communities. Ultimately, our comparative analysis with terrestrial models demonstrates that these marine invertebrates achieve efficient biomass degradation not through microbial expansion, but through extreme functional streamlining and molecular innovation, offering a distinct evolutionary paradigm for marine carbon cycling.

## Introduction

Lignocellulose is the most abundant biopolymer complex on Earth and a major reservoir of fixed carbon ^1,2^. Its deconstruction in nature depends on diverse physical, chemical, and biological mechanisms that have evolved independently across fungi, bacteria, and animals ^2,3^. Terrestrial wood-feeding systems such as termites, passalid beetles, and ruminants rely on dense, taxonomically complex gut microbiomes that convert recalcitrant plant polymers into absorbable metabolites ^4–7^. In contrast, the digestive strategies of marine wood-feeding animals remain comparatively underexplored, despite their pivotal roles in carbon turnover and habitat formation in oceanic wood-fall ecosystems ^1,2^.

Found primarily in shallow marine environments ^8^, shipworms (Bivalvia: Teredinidae) are iconic wood-borers that use wood as both a habitat and a primary food source ^9^. Classical histology and more recent molecular analyses have revealed that shipworms employ a highly unusual digestive strategy: the cecum, where wood is physically processed, is nearly devoid of resident microbiota, whereas dense intracellular bacterial communities are confined to specialized bacteriocytes in the gills^10–12^. These gill endosymbionts, members of the gammaproteobacterial genus *Teredinibacter*, fix nitrogen and secrete cellulolytic enzymes that are transported to the digestive system via the ducts of Deshayes ^13–16^. This “remote digestion” strategy provides the host access to liberated sugars without competition from an endogenous gut microbiota ^14^.

The family Xylophagaidae represents the deep-sea sister lineage to shipworms. These bivalves also consume wood and house bacterial symbionts in their gills ^17,18^, pointing to a single evolutionary origin of xylotrophy in their common ancestor ^19^. However, since diverging, the two families have evolved radically different adult morphologies and occupy entirely distinct oceanic zones ^20^. Although early 16S rRNA surveys showed their symbionts are related ^21^, the evolutionary timeline and genomic stability of this partnership are unknown. This raises a fundamental puzzle: how is such a specialized, spatially segregated digestive system maintained across different environments? It remains to be determined whether shallow-water and deep-sea wood-borers rely on a strictly conserved ancestral blueprint, or if they have evolved independent molecular adaptations to deconstruct lignocellulose.

Efficient lignocellulose degradation typically relies on the synergistic action of highly diverse carbohydrate-active enzyme (CAZyme) repertoires. While terrestrial gut microbiomes achieve this through high taxonomic and functional redundancy ^2^, minimalist symbioses face a distinct evolutionary constraint. One potential molecular adaptation to overcome a streamlined enzymatic toolkit is structural modularity—fusing multiple catalytic domains into single polypeptides to drive localized, synergistic hydrolysis ^22,23^. Such architectural innovations could theoretically compensate for reduced microbial diversity, yet their precise mechanisms and prevalence in marine xylotrophy remain poorly characterized.

Here we use a hologenomic approach—integrating genomes, one of which is resolved to chromosome-level host genomes, metagenomes and metatranscriptomes of gill and gut tissues, and biochemical characterization of key enzymes—to dissect the digestive strategy of a shallow-water shipworm, *Teredo navalis*, and a deep-sea wood-borer, *Xyloredo* sp. We demonstrate that *T. navalis* and *Xyloredo* sp. Holobionts (1) share a conserved genomic framework for gill-based xylotrophy, supporting a single origin of wood feeding and endosymbiosis in Pholadoidea; (2) employ a streamlined, low-redundancy enzymatic strategy for lignocellulose deconstruction that is fundamentally distinct from terrestrial wood-feeding systems such as termites; and (3) have evolved unique molecular innovations—including a lineage-specific dual-domain GH5–GH6 cellulase—that enable highly synergistic cellulose degradation within a compact enzymatic toolkit. By combining microscopy, host and symbiont genomics, comparative CAZyme profiling and functional assays, we propose a unified evolutionary and mechanistic framework for bivalve xylotrophy and expand the known solution space for animal–microbe symbioses involved in biomass decomposition.

## Results and discussion

### *T. navalis* and *Xyloredo* sp. share a conserved anatomical framework for gill-based symbiosis

This study collected the shallow-water shipworm *Teredo navalis* from the Yellow Sea and the deep-sea xylotrophic bivalve *Xyloredo* sp. from the South China Sea (Fig. 1a and Supplymentary Table 1). Consistent with previous observations on teredinid and xylophagaid species^14,17,24,25^, light microscopy and TEM revealed densely packed intracellular bacteria within the gills of both *T. navalis* and *Xyloredo* sp., whereas the cecum contained abundant wood particles but virtually no prokaryotic cells (Fig. 1b,c and Supplementary Fig. 1). FISH with universal bacterial probes confirmed that bacterial signal is strongly localized to gill bacteriocytes (Fig. 1d-f), consistent with previous observations in other xylotrophic bivalves ^14,26^. These observations in representatives of both specimens collected from sunken wood at shallow and bathyal depths support the conclusion that a symbiont-rich gill and a bacteria-poor cecum are not derived traits restricted to particular habitats but instead represent conserved, diagnostic features of xylotrophic bivalves ^19^. These findings also suggest that wood digestion mediated by enzymes produced remotely by gill endosymbionts is a strategy shared by both shallow- and deep-water xylotrophic bivalves and likely predates their ecological divergence. Despite drastic differences in adult physiological organization and habitat — ranging from the extreme elongation of shipworms to the reduced, clam-like *Bauplan* of xylophagaids — the digestive and symbiotic machinery remains evolutionarily conserved. Thus, the distinct morphologies reflect secondary adaptations to depth and substrate availability, belying a unified, ancient origin of symbiotic wood digestion. While the precise anatomical conduits (e.g., ducts of Deshayes) for transporting gill-derived enzymes to the gut in xylophagaids remain to be functionally mapped, the conserved spatial segregation implies an obligate, long-distance molecular trafficking system fundamental to bivalve xylotrophy, as has been demonstrated experimentally in the shipworm, *Bankia setacea* ^16^.

**Fig. 1.**
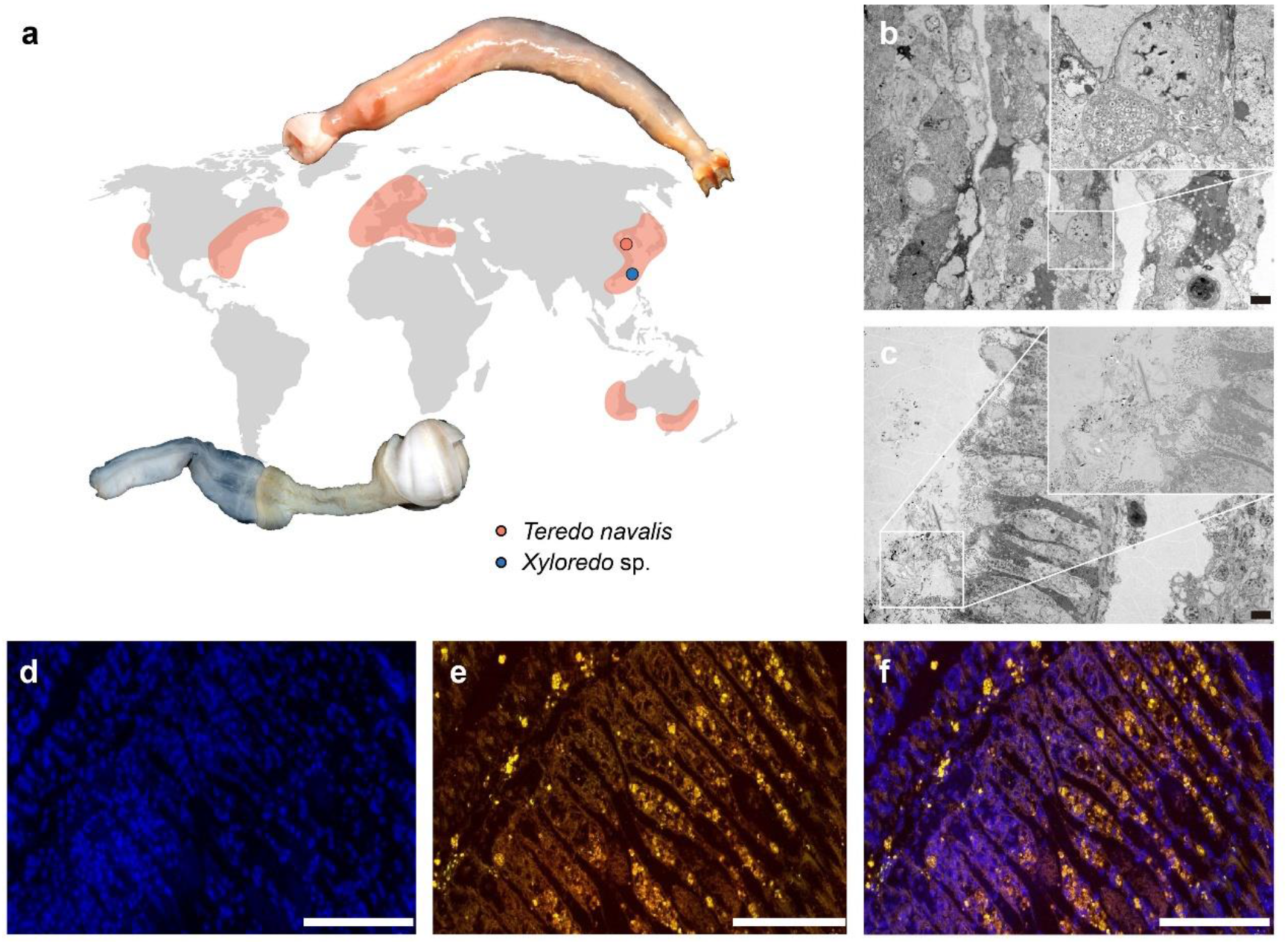
Shallow- and deep-sea xylotrophic bivalves. **a**, The distributions and Bauplan of xylotropic bivalves. *Teredo navalis*, depicted at the top, is a globally distributed shallow-water species. A specimen was collected from the intertidal zone of the Yellow Sea for this study. *Xyloredo* sp., shown at the bottom, was collected from the Site F cold seep in the South China Sea at a depth of 1100 m. The orange-shaded area on the map indicates the documented distribution of *T. navalis*, while the blue dot indicates the sampling site of *Xyloredo* sp. **b**, A transmission electron micrograph illustrating endosymbionts residing within the bacteriocytes of shipworm gills. **c**, In contrast, the cecum was almost devoid of bacteria (Scale bars: b and c, 5 μm). **d–f**, FISH images showing (d) nucleic acid stain (DAPI, blue), (e) *Teredinibacter* probe (Cy3, yellow), and (f) merged images (Scale bars: d–f, 200 μm).

### Host genomes reveal conserved chromosomal evolution and lineage-specific expansions enabling lignocellulose digestion

We generated a chromosome-level assembly for the *T. navalis* genome (∼914 Mb; 19 chromosomes) and a highly contiguous assembly for *Xyloredo* sp. (∼644 Mb), both with BUSCO completeness exceeding 92% (Extended Data Fig. 1 and Supplementary Fig. 2 and Supplymentary Tables 2-4). Macrosynteny analyses indicated that both genomes retain an ancestral molluscan chromosomal architecture, exhibiting relatively few large-scale rearrangements (Fig. 2 a,b). A notable shared feature is the fusion of ancestral molluscan linkage groups (MLGs) 16 and 18 into a single chromosome in *T. navalis* (Fig. 2a), with orthologous fused contigs observed in *Xyloredo* sp. (Fig. 2b). However, this chromosomal fusion is conserved across members of the clade Imparidentia and Polyplacophora, including the closest sequenced relatives of these xylotrophs—the soft-shell clam (*Mya arenaria*) and the zebra mussel (*Dreissena polymorpha*) ^27^(Extended Data Fig. 2). Consequently, the radical morphological divergence observed in xylotrophic bivalves is not accompanied by extensive syntenic rearrangements.

**Fig. 2.**
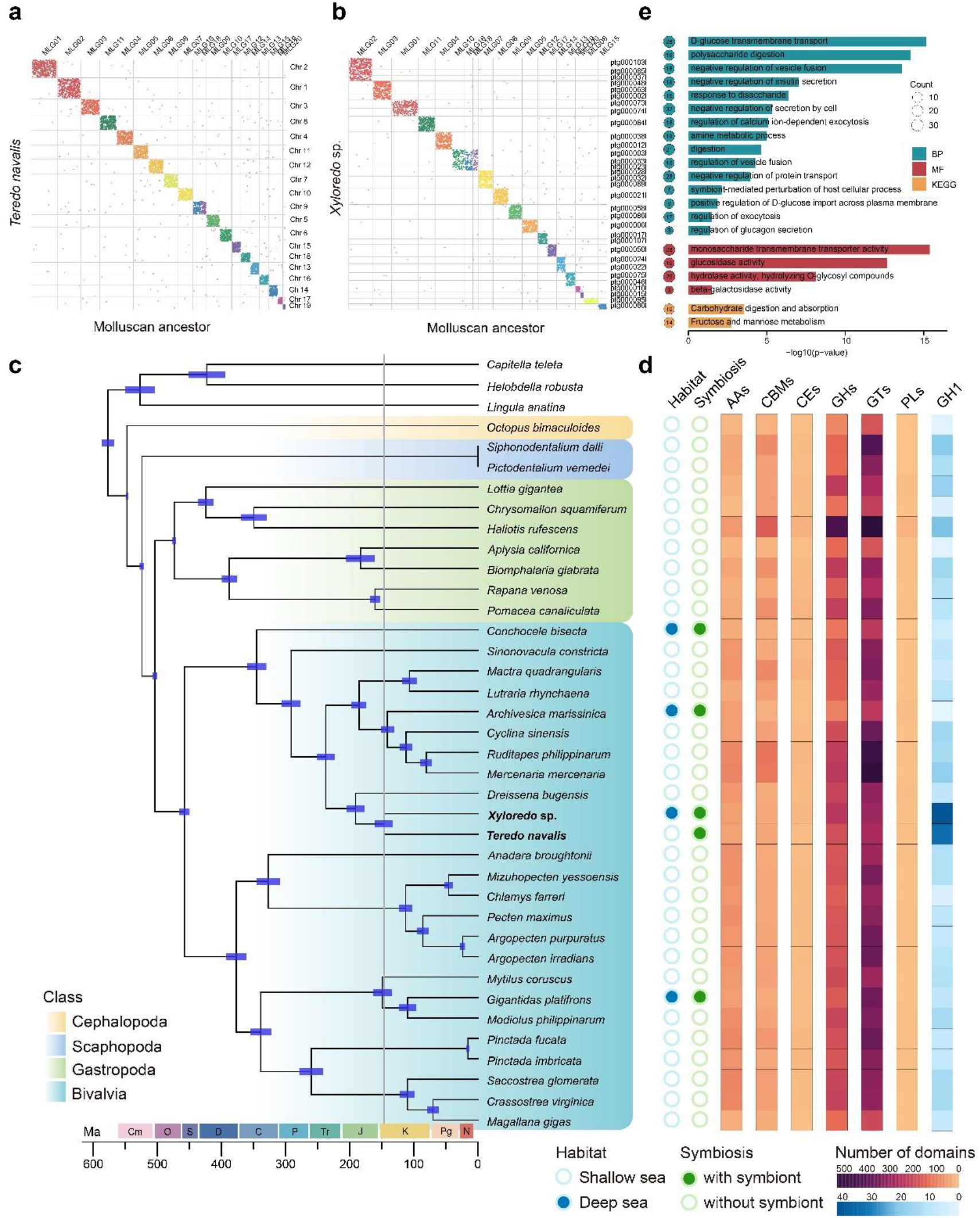
Evolution and landscape of endogenous carbohydrate-active enzymes in xylotrophic bivalves and their relatives. Oxford plots displaying conserved synteny between the ancestral molluscan linkage groups (MLGs; horizontal axes) and the genomes of *T. navalis* (**a**) and *Xyloredo* sp. (**b**) (vertical axes). For panel **b**, only contigs larger than 10 Mb are shown (see Supplementary Fig. 3 for the complete plot). **c**, Phylogenomic position of xylotropic bivalves. The phylogenetic tree was calibrated at six nodes (see Supplementary Fig. 4) using fossil data to estimate divergence times. The estimated divergence times are represented on the branches of the tree, with the 95% confidence intervals indicated by blue bars. For clarity, the splitting time of *T. navalis* and *Xyloredo* sp. is marked by a vertical grey line. **d**, Heat map of carbohydrate-active enzyme-related domains: Auxiliary Activities (AAs), carbohydrate-binding modules (CBMs), carbohydrate esterases (CEs), glycoside hydrolases (GHs), glycosyltransferases (GTs), and polysaccharide lyases (PLs). The circles represent the habitats and symbiosis of different species. **e**, Gene Ontology (GO) and KEGG pathway enrichment for expanded gene families of the last common ancestor of xylotrophic bivalves. The enriched terms, with corrected p-values < 0.05 (calculated using the BH method), are presented, and the size of the circles corresponds to the number of expanded genes for each term. The complete GO and KEGG enrichment analysis results are provided in Supplementary Tables 5 and 6, respectively. BP, biological process; MF, molecular function; Cm, Cambrian; O, Ordovician; S, Silurian; D, Devonian; C, Carboniferous; P, Permian; Tr, Triassic; J, Jurassic; K, Cretaceous; Pg, Paleogene; N, Neogene.

Morphological adaptations across Bivalvia, particularly trait reductions associated with specialized lifestyles, are often underpinned by modifications in conserved developmental gene networks. For instance, the loss of the homeobox gene Antennapedia (*Antp*) has previously been associated with the loss of byssus secretion in oysters^28^. To broaden our understanding of homeobox gene evolution across Bivalvia, we investigated the structural organization of Hox and ParaHox clusters in xylotrophic bivalves (Extended Data Fig. 3). A conspicuous absence of the *Antp* gene is observed across disparate lineages, ranging from oysters within Pteriomorphia to xylotrophs and their close relatives within Heterodonta (Extended Data Fig. 3). This scattered distribution indicates that *Antp* has been lost independently multiple times throughout bivalve evolution. Similar to oysters, xylotrophic bivalves and their relatives (e.g., *Mya arenaria*) lack an adult byssus. However, notable counterexamples suggest that the retention of the *Antp* gene and adult byssus formation are evolutionarily decoupled in certain lineages: *Ruditapes philippinarum* retains *Antp* but does not form an adult byssus, whereas *Dreissena polymorpha* develops a robust adult byssus despite lacking *Antp* (Extended Data Fig. 3). Given that the byssus is a foot-derived secretion^29^, we hypothesize that the recurrent loss of *Antp* is more fundamentally linked to the evolutionary reduction or complete loss of the foot itself. In the context of xylotrophic evolution, foot reduction and specialization are direct consequences of their transition to a wood-boring lifestyle, where the foot serves not as an organ of locomotion but rather as an anchor stabilizing the animal within the burrow and countering the forces generated by the adductor muscles and valves during boring.

Unlike terrestrial xylophagy, which evolved independently across multiple insect lineages ^2^, marine bivalve xylotrophy arose from a single evolutionary origin (Fig. 2c). Molecular clock estimates date the divergence of the shallow-water Teredinidae and deep-sea Xylophagaidae to approximately 147 million years ago (Fig. 2c and Supplementary Fig. 4), a timeline supported by the minimal Cenomanian divergence implied by fossil evidence^30^. Following this ancient split, the clade underwent profound ecological radiation: Teredinidae adapted primarily to warm, shallow waters, whereas Xylophagaidae colonized perennially cold environments ranging from sublittoral zones to the deep-sea.

Despite exceeding 100 million years of extreme ecological divergence, these two lineages retain a highly conserved repertoire of all six major CAZyme classes (AAs, CBMs, CEs, GHs, GTs, and PLs)—a pattern consistent across molluscs, independent of habitat or symbiotic status (Fig. 2d). Strikingly, family-level profiling revealed parallel expansions of glycoside hydrolase families 1 (GH1) in the two xylotrophic bivalves *T. navalis* and *Xyloredo* sp., contrasting with patterns observed in non-wood-feeding relatives (Fig. 2d). While multi-domain GH1 enzymes are widely distributed across various animal lineages, the unique six-module GH1 architecture identified in the shipworm *Lyrodus pedicellatus* was previously characterized as a specialized adaptation for wood digestion^24^. Here, we report similar multi-modular GH1 configurations within the genomes of *T. navalis* and *Xyloredo* sp. (Supplementary Fig. 5). This conserved structural expansion strongly suggests that such specific multi-modularization is not a lineage-specific anomaly, but rather a broader, shared evolutionary adaptation fundamental to the xylotrophic bivalves.

Gene families expanded in the last common ancestor of these two bivalve lineages exhibit significant Gene Ontology (GO) enrichment in processes related to host-symbiont interactions, vesicle trafficking, and exocytosis (Fig. 2e). These expansions dovetail with the complex regulation required for the host–microbe interface and secretory digestion^31^. Notably, among the ancestrally expanded genes, those encoding multi-domain GH1 are significantly enriched for polysaccharide digestion (GO:0044245), glucosidase activity (GO:0015926), response to disaccharide (GO:0034285), and digestion (GO:0007586), and are mapped to the carbohydrate digestion and absorption KEGG pathway (ko04973). Furthermore, expanded GH5 genes are implicated in mannose metabolism (ko00051), while genes encoding sodium-dependent glucose transporters (*MFSD4*) are highly enriched for monosaccharide transmembrane transporter activity (GO:0015145) and D-glucose transmembrane transport (GO:1904659) (Fig. 2e). Collectively, these lines of evidence demonstrate that their common ancestor had already established the robust capacity for the degradation, efficient absorption, and utilization of wood-derived sugars.

### Gill microbiomes are dominated by deeply branching *Teredinibacter* lineages with expanded lignocellulolytic potential

Metagenomic analyses reveal that the gills of both *T. navalis* and *Xyloredo* sp. harbor a high microbial abundance (nearly 50%), heavily dominated by *Cellvibrionaceae*, whereas their caeca are virtually devoid of bacteria (Fig. 3a and Supplementary Tables 7 and 8). This striking pattern of microbial compartmentalization is consistent with previous microscopy-based reports for Teredinidae and Xylophagaidae^14,17^. Metagenomic binning successfully recovered near-complete metagenome-assembled genomes (MAGs; >96% completeness) representing the dominant gill symbiont within each host (Supplementary Tables 7 and 8). Notably, these endosymbionts retain large, non-reduced genomes (4.3–5.6 Mb, 45.9%–49.8% G+C) (Fig. 3b and Supplementary Tables 7 and 8 and Supplementary Fig. 6). This stands in stark contrast to the highly reduced, AT-rich genomes typical of many obligate insect endosymbionts (<710 kb, mostly 13.5% – 33.2% G+C)^32,33^, but aligns with the previously sequenced genome of *Teredinibacter turnerae* T7901 (5.2 Mb, 50.8% G+C)^13,15^. Specifically, the examined *T. navalis* specimen harbored a remarkably simple symbiont community comprising just two detectable species, consistent with previous observations of one to five symbiont species in other Teredinidae^12,14,34^. Similarly, a single symbiont species dominated the microbial community of the gills of this specimen of *Xyloredo* sp. (accounting for approximately 56.06% of the total microbiome), representing, to our knowledge, the first assessment of gill symbiont community complexity within the Xylophagaidae. Average amino acid identity (AAI) and phylogenomic reconstructions classify both MAGs within the genus *Teredinibacter* (*Cellvibrionaceae*, Gammaproteobacteria), though they form early-diverging lineages relative to the cultured model *T. turnerae* and other described strains(Fig. 3c and Supplementary Fig. 7 and 8 and Supplementary Table 9).

**Fig. 3.**
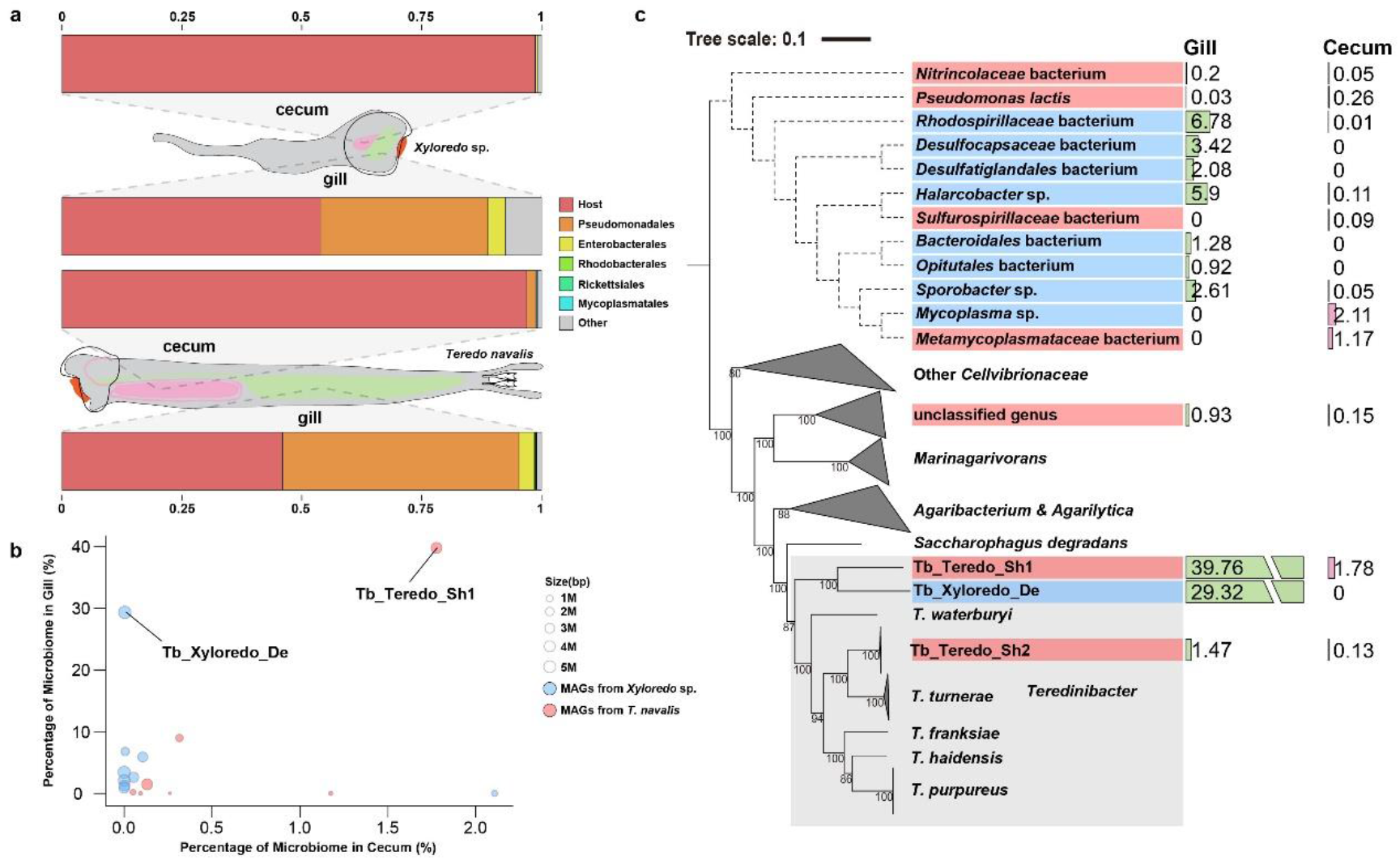
Xylotrophic bivalves exhibit a clear compartmentalization of microbial biomass. **a**, Relative abundance of bacterial orders in the gills and cecum of *Xyloredo* sp. (upper panel) and *T. navalis* (lower panel). Each color represents a specific bacterial order, and the label “Other” denotes orders not shared between the two species. **b**, Metagenomic binning of the gills and cecum of xylotrophic bivalves, along with the relative abundance of microbial communities in these tissues. Blue circles represent Metagenome-assembled genomes (MAGs) from *Xyloredo* sp., while red circles represent MAGs from *T. navalis*. The size of the circles corresponds to the genome sizes of the MAGs. **c**, Phylogenetic tree of *Cellvibrionaceae*, with MAGs not belonging to *Cellvibrionaceae* selected as the outgroup. Bootstrap support values greater than 70% are indicated. MAGs are labeled according to the color scheme used in (**b**), and the genus *Teredinibacter* is highlighted with a gray frame. The bars on the right display the relative abundances of each MAG in the gills and cecum.

Functionally, while one of the symbiont species within *T. navalis* (Tb_Teredo_Sh2) possesses nitrogen-fixation (*nif*) operons—similar to *T. turnerae* T7901^13^(Supplementary Table 10)—the dominant symbiont of *T. navalis* and *Xyloredo* sp. entirely lack all classes of *nif* genes(Supplementary Table 10). Instead, they possess a complete pathway for dissimilatory nitrate reduction to ammonium (DNRA). Beyond nitrogen metabolism, all symbionts of *T. navalis* and *Xyloredo* sp. retain key components of the Sec pathway (*secAYEG*) alongside complete Type II (T2SS) and Type V (T5aSS and T5bSS) secretion systems(Supplementary Table 11). Yet, they conspicuously lack the Type III and IV systems typically associated with pathogenic or tightly host-associated bacteria, mirroring previous observations in *T. turnerae* T7901^13^(Supplementary Table 11). Additionally, we annotated a complete Type I (T1SS) and a nearly complete Type VI (T6SS) secretion system within the symbionts of both bivalve species (Supplementary Table 11 and Supplementary Fig. 9). This distinctive genomic profile points toward a functional strategy heavily centered on the robust secretion of extracellular hydrolases, without reliance on type III or Type IV secretion of effector that establish or maintain host interactions in other intracellular symbionts. Furthermore, given that some shipworm symbionts are known to be capable of independent survival outside their hosts ^13,35^, these combined genomic and physiological traits strongly suggest that these microbes are facultative, rather than obligate, endosymbionts.

These *Teredinibacter* lineages showed large complements of CAZymes, including expanded GH5 and GH9 families, exceeding those found in free-living *Cellvibrionaceae* relatives (Extended Data Fig. 4a and Supplementary Tables 12 and 13). A substantial proportion of these enzymes, 32% and 71%, respectively, are characterized by the presence of CBMs and signal peptides (Extended Data Fig. 4 b,c and Supplementary Table 14), consistent with a lifestyle that couples intracellular residence with extensive enzyme secretion. The dominance of *Teredinibacter* in the gills of both Teredinidae and Xylophagidae, combined with their basal phylogenetic positions, supports a phylogenetically stable, long-term association and reinforces the hypothesis of a single origin of xylotrophic endosymbiosis in bivalves^19^.

### Host and symbionts jointly orchestrate a three-component holobiont pathway for lignocellulose degradation

By integrating metagenomic, metatranscriptomic, and domain-based annotations, we constructed a holobiont-level model of lignocellulose deconstruction across both species, comprising three components: lignin modification, hemicellulose degradation, and cellulose degradation (Fig. 4a). The model suggests a highly coordinated division of labor between the host and its symbionts (Fig. 4b-h). Host-derived (endogenous) lignin-modifying enzymes (LMEs)-such as laccases, peroxidases, and auxiliary oxidoreductases-account for an average of approximately 89% of the total contribution, indicating that lignin modification is primarily driven by the host^36^ (Fig. 4b). This echoes the oxidative strategies observed in fungal lignin degradation^37^, but contrasts sharply with soil isopods, where symbionts dominate this process^38^. For hemicellulose degradation, endogenous enzymes provide approximately 29% of the identified domains, whereas the symbionts exclusively supply the remaining 71%. Specifically, the host genomes encode enzymes from the GH5, GH10, GH8, and GH29 families to facilitate hemicellulose debranching, complementing the GH5 and GH10 repertoire derived from the symbionts (Fig. 4c). Furthermore, a distinct division of labor is evident regarding the endo- and exo-cleavage of hemicellulose. For instance, the host’s contribution is restricted to the GH27, GH2, GH38, and GH47 families, whereas the symbiont genomes encode a wider array of hemicellulases (Fig. 4d,e). Indeed, the number of hemicellulase families encoded by the symbionts is nearly 1.5 times that of the host—a trend particularly pronounced in *T. navalis*. This underscores the predominant role of the microbiota in hemicellulose degradation, a dynamic analogous to that observed in the terrestrial isopod *A. vulgare* ^38^. The cellulose degradation component, overall, indicates a synergistic mechanism between the host and its symbionts, with each partner contributing approximately 50% to the degradative process (Fig. 4f-h). Notably, in our study, this specific catalytic function involved exclusively bacteria-derived exoglucanases (GH6) (Fig. 4g), consistent with previous observations in the shipworms *B. setacea* and *L. pedicellatus* ^14,24^. Because this distinct enzymatic activity is absent in several terrestrial xylotrophic hosts^38–40^, it appears that the symbionts have not merely supplemented the host’s digestive repertoire, but have evolved highly specialized machinery for core cellulose depolymerization, representing a profound evolutionary integration within the holobiont.

**Fig. 4.**
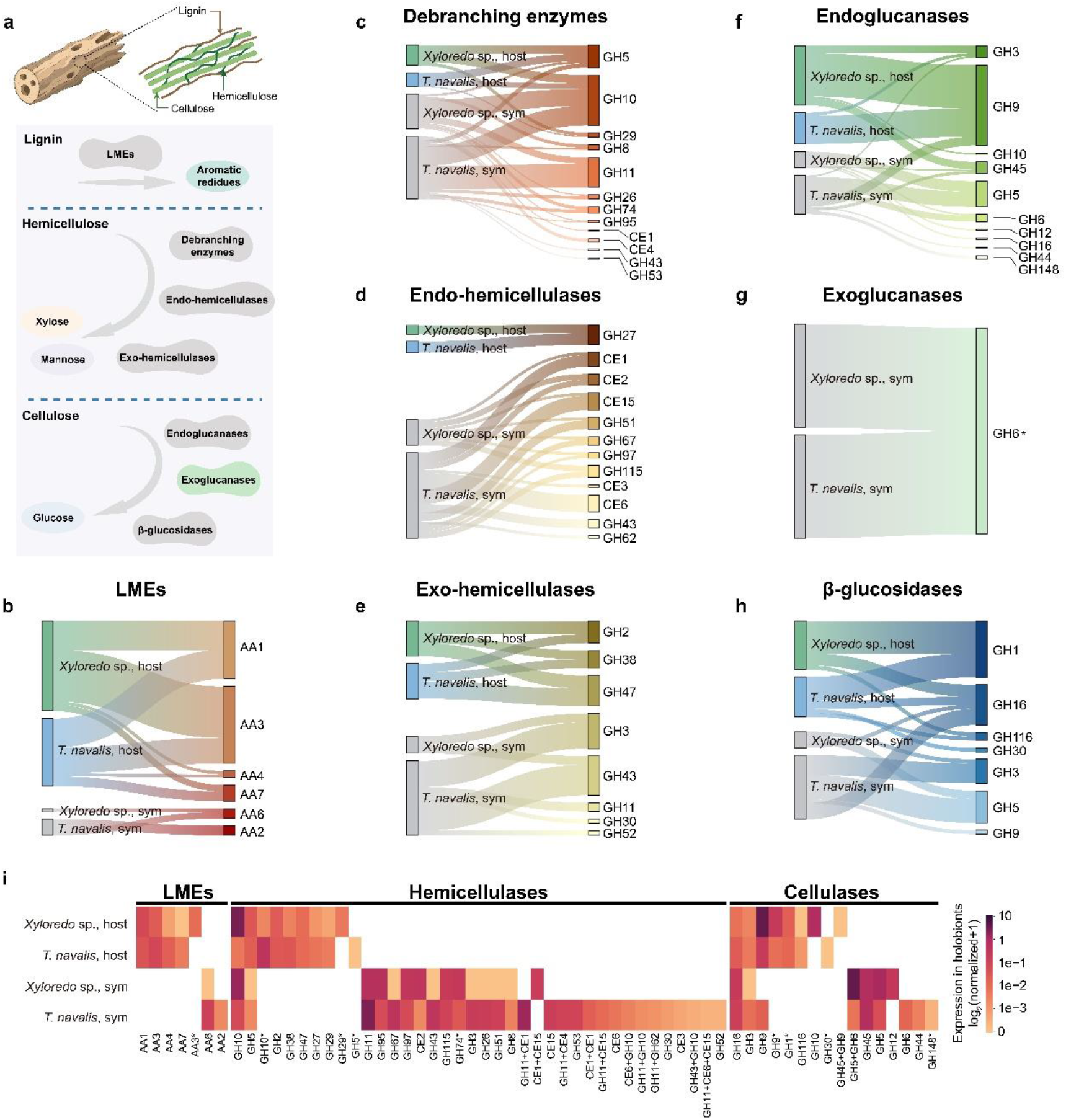
Potential contribution of host and symbiont genomes to lignocellulose degradation in xylotrophic bivalve holobionts. **a**, Diagrammatic representation of the enzymatic processes that may contribute to lignocellulose degradation in the caeca of xylotrophic bivalves. The cecum serves as the principal site for wood digestion, playing a key role in the breakdown of lignin, hemicellulose, and cellulose. Hemicelluloses are branched polymers composed of various sugar monomers; for clarity, only the predominant components, xylose and mannose, are presented here. **b–h**, Contributions of the host and symbiont genomes (left) to the total repertoire of lignocellulose-active CAZymes by family (right) in the respective hologenomes of *T. navalis* and *Xyloredo* sp. by function: (**b**) Lignin-modifying enzymes (LMEs) that modify lignin for its breakdown; (**c**–**e**) Hemicellulases, which include: (**c**) Debranching enzymes responsible for cleaving side chains, (**d**) Endo-hemicellulases that mediate internal scission of the primary chain, and (**e**) Exo-hemicellulases that facilitate the release of monomeric sugars. (**f**–**h**) Three types of cellulases: (**f**) Endoglucanases, (**g**) Exoglucanases, and (**h**) β-Glucosidases. An asterisk in (**g**) indicates that this domain is derived from a GH5–GH6 dual-domain enzyme. Connecting lines indicate occurrence of CAZyme families in each host genome or metagenome. Line thickness is proportional to the number of domains. Scales are adjusted independently for each panel and are not directly comparable; exact domain counts are provided in Supplementary Table 18. **i**, Relative expression of lignocellulose-degrading genes in xylotrophic holobionts. Gene entries containing multiple catalytic activity domains from the same gene family are marked with an asterisk for clarity.

Supporting this model, host transcripts dominate the lignin modification pathway (accounting for an average of ∼80% of total expression), whereas symbiont-derived transcripts predominantly drive hemicellulose (71.50%) and cellulose (75.53%) degradation (Fig. 4i and Supplementary Table 15). Remarkably, this three-component model - host-driven lignin oxidation, symbiont-dominated hemicellulose and cellulose depolymerization - was conserved across *T. navalis* and *Xyloredo* sp., as demonstrated by their highly congruent gene expression profiles and consistent division of labor between the host and symbionts (Fig. 4i). These findings further bolster the deep evolutionary foundations of this biochemical mechanism, providing strong evidence for the synchronized metabolic functions of the host and its symbionts in lignocellulose breakdown.

Previous proteomics-based studies reported conflicting host-symbiont enzyme contributions in the cecum, inherently capturing only physiological snapshots^14,24^. By characterizing the genomic arsenals of *T. navalis* and *Xyloredo* sp., we instead define the holobiont’s fundamental coding potential, revealing a deeply conserved and complementary lignocellulolytic gene repertoire. Reconciling this genomic blueprint with prior protein data, we speculate that the relative contribution of host and symbiont enzymes is not a fixed attribute but a dynamically modulated physiological process that adapts to the holobiont’s environment and metabolic demands.

### Beyond termites: a minimalist but synergistic digestive strategy

To place xylotrophic bivalves in a broader evolutionary and ecological context, we compared the lignocellulase repertoires of *T. navalis* and *Xyloredo* sp. with those of representative higher and lower termites, which rank among the most efficient terrestrial lignocellulose degraders ^7,39,41,42^ (Fig. 5 and Supplementary Table 16). This comparison highlights a starkly distinctive digestive strategy in the bivalves. Termite hindguts harbor densely packed, open microbial ecosystems—comprising hundreds of species—that collectively produce a vast but functionally overlapping array of lignocellulases ^39,43^. In contrast, our metagenomic analyses reveal that bivalve hosts harbor remarkably simple symbiont communities, in this case consisting of only two dominant symbiont species in *T. navalis* and a single species in the gills of *Xyloredo* sp. (Fig. 3) Consequently, the total number of lignocellulose-active domains in termite systems is estimated to be two to five times higher than in the bivalves (Fig. 5a). For instance, a termite hindgut contains an average of ∼121 endo-cellulase domains, compared to only 44 and 50 observed in *T. navalis* and *Xyloredo* sp., respectively (Supplementary Table 17). Despite this reduced domain pool, bivalve symbioses maintain broad coverage of lignocellulose-degrading functions while exhibiting less than 50% of the functional and family-level redundancy observed in termites (Fig. 5b,c and Extended Data Fig. 5 and Supplementary Tables 17 and 18). This lower redundancy stems fundamentally from the compositional simplicity of the bivalve symbiont community, in contrast to multi-taxa termite guts, where diverse microbes maintain parallel solutions to the same catalytic challenges. At the family level, termite symbionts act as the primary reservoir of redundancy (e.g., GH4, GH55, and GH158). While this distinguishes them from bivalve symbionts, certain families (e.g., GH52, CE3, GH148) remain exclusive to the bivalves (Fig. 5d). The bivalve hosts and their symbionts encode a significantly higher proportion of multi-domain carbohydrate-active enzymes (CAZymes) (Fig. 5e), potentially compensating for a reduced number of gene families through structural modularity. A prime example of this evolutionary strategy is a symbiont-encoded, bi-catalytic GH5 – GH6 enzyme responsible for the synergistic degradation of cellulose (Figs. 6 and 7).

**Fig. 5.**
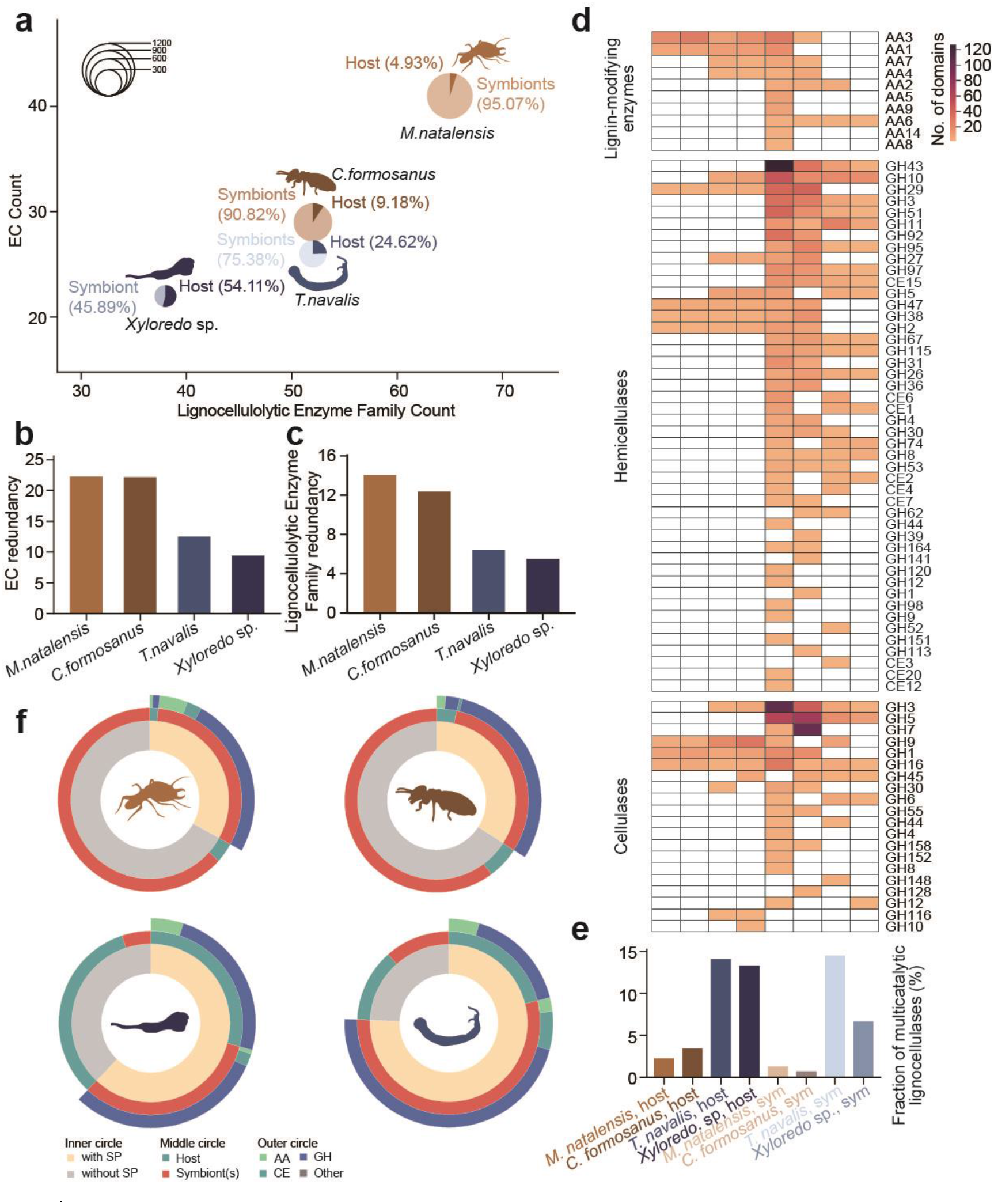
Comparison of lignocellulolytic enzyme systems in xylotrophic bivalves and termites. **a**, Domain-level landscape of lignocellulases in *Macrotermes natalensis*, *Coptotermes formosanus*, *T. navalis*, and *Xyloredo* sp. The axes represent the number of lignocellulolytic enzyme families (*x*-axis) and their associated functional EC categories (*y*-axis) for each symbiosis. The pie charts illustrate the relative contributions of the host and symbionts to lignocellulase activity. Chart size is proportional to the total number of lignocellulase domains in each system: *M. natalensis* (913), *C. formosanus* (643), *T. navalis* (325), and *Xyloredo* sp. (207). Redundancy indices of lignocellulases based on Enzyme Commission (EC) categories (**b**) and gene families (**c**) across the four symbioses. Redundancy is defined as the average number of lignocellulase domains per EC category or gene family within each system. **d**, Domain-level classification of lignocellulases into lignin-modifying enzymes, hemicellulases, and cellulases across the four symbioses. Termites exhibit a higher abundance of these enzyme families (*M. natalensis*: 56, *C. formosanus*: 44) compared to xylotrophic bivalves (*T. navalis*: 42, *Xyloredo* sp.: 33). Functional contributions across the holobionts show a similar pattern (Extended Data Fig. 5 and Supplementary Table 17). **e**, Proportion of genes encoding multi-catalytic domains among all lignocellulase genes in xylotrophic bivalves and termites. **f**, Signal peptide profiles of lignocellulases in xylotrophic bivalves and termites. Concentric circles (from inner to outer) display the proportion of lignocellulases possessing signal peptides, their origin (host or symbiont), and the associated carbohydrate enzyme class.

**Fig. 6.**
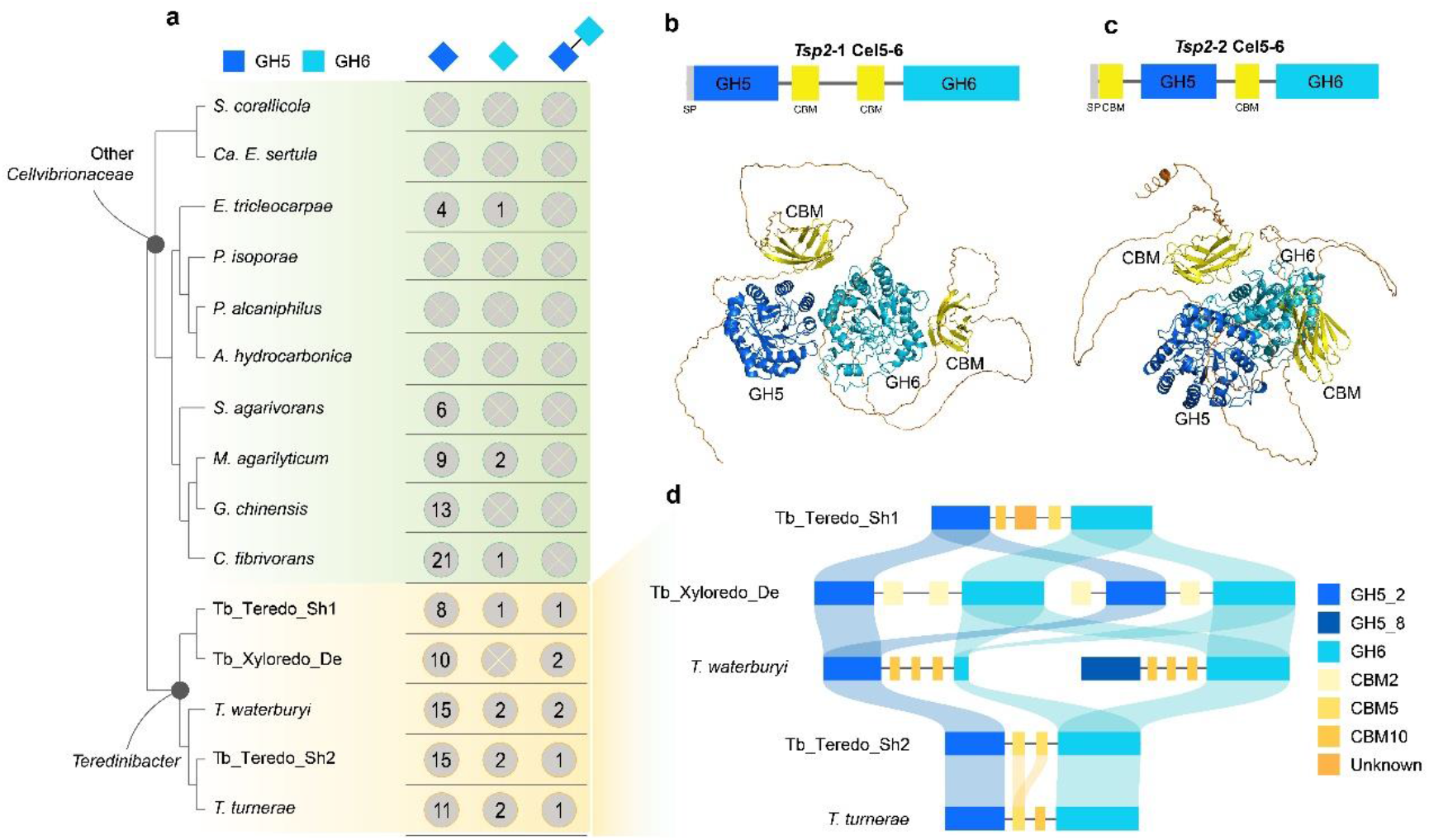
T*e*redinibacter-specific bi-domain GH5–GH6 enzyme. **a**, Phylogenetic distribution of GH5-only, GH6-only, and bi-catalytic GH5–GH6 enzymes within *Cellvibrionaceae*. Colored boxes represent stylized domain architectures. The *Xyloredo* sp. symbiont (Tb_Xyloredo_De) encodes two bi-domain GH5–GH6 enzymes (**b**, **c**). One of these is *Tsp2-1* Cel5-6 (**b**), which starts at the N-terminal with a signal peptide (SP), followed by a glycosyl hydrolase 5 (GH5) domain, two cellulose-binding modules (CBMs), and a glycosyl hydrolase 6 (GH6) domain at the C-terminal. The other enzyme exhibits an inversion in its domain structure (*Tsp2-2* Cel5-6, **c**). The unstructured linkers, CBMs, and GH5 and GH6 catalytic modules are colored gray, yellow, blue, and cyan, respectively. Protein structure predictions were conducted using AlphaFold3 (https://alphafoldserver.com/). **d**, Ribbon diagram illustrating the homology relationships among domains in the bi-domain GH5–GH6 enzymes of *Teredinibacter*.

Beyond genomic redundancy, xylotrophic bivalves and termites exhibit fundamentally divergent enzyme localization and secretion strategies (Fig. 5f). In termites, microbial lignocellulose degradation is largely internal or contact-dependent; many cellulases operate within protist cells or are scavenged by surface-adhering bacteria, yielding sugars that undergo microbial fermentation before becoming accessible to the host ^2,39^. Consequently, a meta-analysis indicates that only ∼14–26% of termite hindgut CAZymes carry signal peptides (Extended Data Fig. 6). In stark contrast, the bivalve holobiont is heavily focused on secretion. We found that ∼60% of CAZymes in xylotrophic bivalves possess secretory signals, representing a pronounced enrichment, particularly among glycoside hydrolases (GHs) (Fig. 5f and Extended Data Fig. 6). This outright solubilization of cellulose into glucose facilitates the direct utilization of simple sugars by both the host and its symbionts. Furthermore, while only Sec signal peptides (Sec/SPI) were predicted in bivalve host enzymes, their symbionts utilize a broader array of export mechanisms, including lipoprotein signal peptides (Sec/SPII, ∼15–20%) and, notably in deep-sea symbionts, Tat signal peptides (Tat/SPI, ∼15%) (Extended Data Fig. 6). Collectively, this massive investment in secretory pathways reflects an adaptation for the long-distance trafficking of enzymes from the gill to the cecum. While the termite host contributes to digestion via salivary GH9 cellulases produced in the foregut ^44^, the endogenous enzymes of xylotrophic bivalves are secreted by the digestive gland directly into the cecum ^24^. Thus, rather than relying on an open gut community to locally digest wood particles, xylotrophic bivalves spatially separate enzyme production from degradation, transporting a customized suite of host-and symbiont-derived enzymes into a dedicated digestive chamber.

In the broader spectrum of symbioses, these bivalves and insects represent two functional extremes of host-microbe integration. Termites, alongside intermediates like ruminants, outsource digestion to a complex, largely anaerobic gut microbiome, relying heavily on internal microbial fermentation ^45^. In this model, resident microbes consume a significant portion of the carbohydrates for their own metabolism, providing the host primarily with fermentation by-products such as short-chain fatty acids ^2,39^. Conversely, xylotrophic bivalves internalize an microaerobic microbial community within their gills ^46^. By exporting enzymes without requiring symbiont migration, the cecum of xylotrophic bivalves functions analogously to a cell-free industrial bioreactor, optimized for the complete enzymatic hydrolysis of lignocellulose into monomers (e.g., glucose) ^24^. The absence of a large resident microbial community in the gut eliminates competition for these liberated sugars, allowing the host to directly absorb simple carbohydrates. Therefore, while termites have converged upon an internal fermentation strategy, xylotrophic bivalves employ a compartmentalized enzymatic digestion model—one conceptually convergent with the external digestion strategies of free-living fungi. The evolutionary success of this shipworm strategy is evident in their global distribution and fossil record^8,30,47^; it emphasizes that even in nutrient-poor marine realms, a tight host-symbiont collaboration can unlock vast energy resources (sunken wood), profoundly impacting carbon turnover in the oceans.

### A lineage-specific GH5–GH6 fusion as a key molecular innovation

Initially identified in *T. turnerae* T8201^48^, the GH5–GH6 fusion unites an endoglucanase (GH5) and an exocellulase (GH6) into a single polypeptide^14,49^. Our work now provides an integrative view, elucidating the evolutionary trajectory, spatial architecture, and biochemical mechanism of the full-length enzyme, thereby resolving a longstanding gap in the understanding of this unusual biocatalyst. Within the symbiont CAZyme repertoire, we identified a dual-domain cellulase (GH5–GH6) exclusively encoded by bivalve-associated *Teredinibacter* lineages (Fig. 6a and Supplementary Figs. 10 and 11). These GH5–GH6 catalytic modules are typically separated by two or three carbohydrate-binding module (CBM) domains (Fig. 6b-d). In *T. navalis* symbiont metagenomes, these CBMs are interconnected by poly-serine linkers, whereas in proteins derived from deep-sea species, they are joined by multi-repeat sequences such as TTSGS or TTSSS. These linkers are predicted to act as flexible, disordered spacers^50,51^. Structural alignments of the GH5 and GH6 domains across *Teredinibacter* dual-domain enzymes reveal highly conserved 3D architectures (mean TM-scores > 0.9; mean LDDT > 0.8), despite variations in their primary sequences (Extended Data Fig. 7). In contrast, the intervening CBM domains have undergone frequent shuffling (Fig. 6d and Extended Data Fig. 8), indicating strong evolutionary conservation of the core catalytic machinery paired with rapid adaptation for substrate recognition — even if its constituent catalytic domains likely arose via multiple ancient, independent fusion events (Extended Data Fig. 8b). Such a multifunctional architecture is exceedingly rare in termite or fungal systems, which instead rely on multi-enzyme complexes to achieve endo–exo synergy ^52,53^.

Modular cellulases typically tether catalytic domains to CBMs rather than directly to other catalytic modules ^54^. However, structural modeling demonstrates that the GH5 and GH6 domains fold to form overlapping catalytic clefts, bringing their respective activities into extreme spatial proximity (Fig. 7a and Supplementary Figs. 12 and 13). Crucially, certain residues physically span both domains, creating an extended, shared substrate-binding groove that is highly atypical among CAZymes (Fig. 7b). Whereas conventional cellulase synergy relies on random molecular collisions or multi-enzyme complexes, molecular docking reveals that this dual-domain architecture physically couples endo-cleavage with processive exo-cleavage (Fig. 7c,d). Furthermore, both domains exhibit critical subsite extensions: the GH5_2 cleft features putative +3 and +4 subsites, which are complemented by a +4 subsite within the adjacent GH6 cleft (Fig. 7c,d). This fusion provides the structural basis for the accelerated, concerted degradation of long crystalline cellulose chains. From an evolutionary perspective, this represents a striking molecular compensation: achieving the catalytic synergy of a complex microbial consortium within a single polypeptide.

**Fig. 7.**
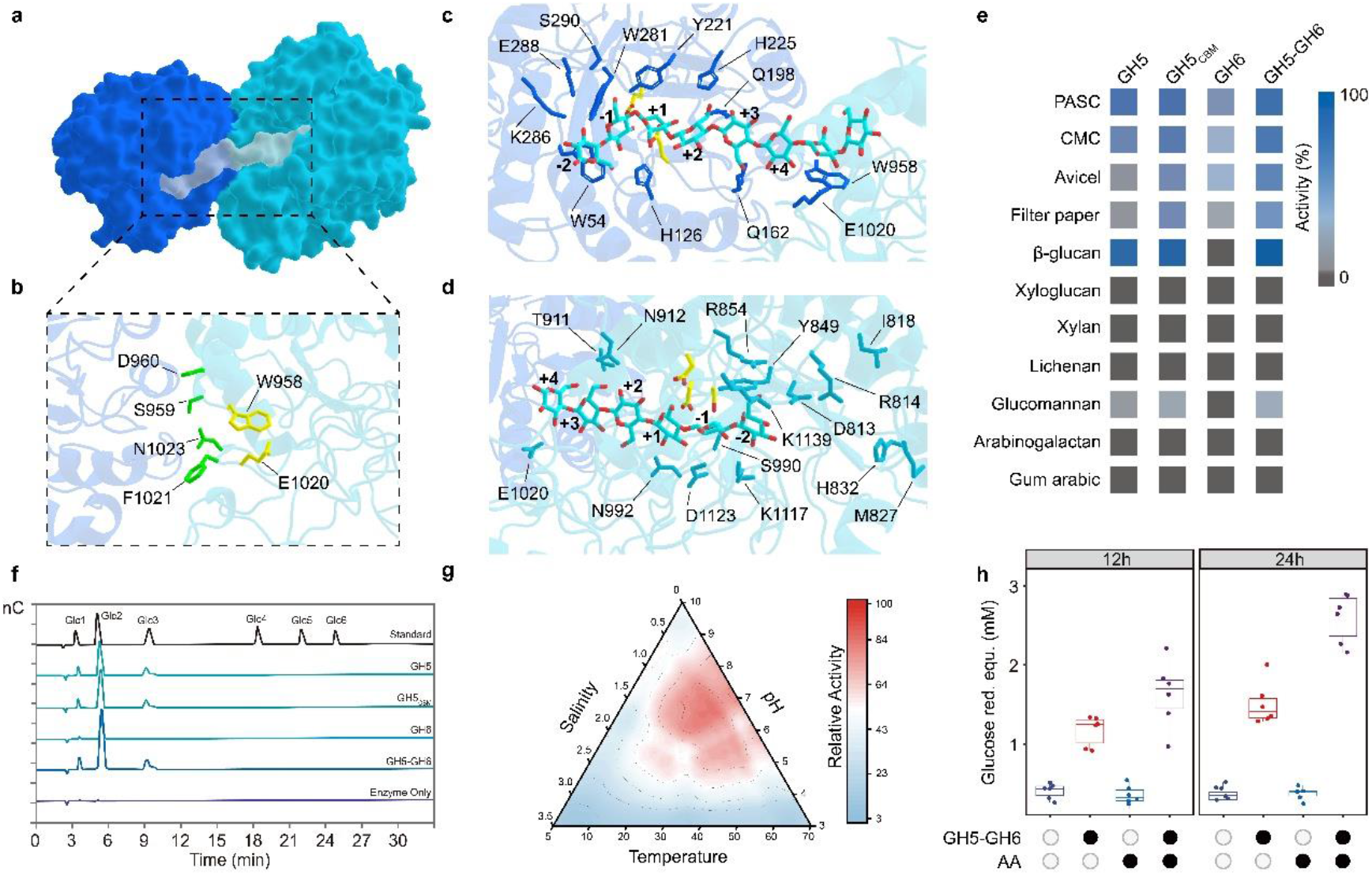
Intramolecular synergy mechanism for cellulose degradation within the dual-domain GH5-GH6 enzyme. For clarity, carbohydrate-binding modules (CBMs) are omitted from the structural representations. **a**, The dual-domain GH5–GH6 enzyme adopts a compact architecture in which the endoglucanase (GH5) and exocellulase (GH6) domains are closely juxtaposed, forming a novel binding pocket, as represented on the white surface of cellooctaose. The surfaces of the GH5 and GH6 domains are rendered in blue and cyan, respectively. **b**, The GH5 and GH6 binding pockets share residues at the inter-domain interface. Six side chains shown within the GH5 pocket are contributed by the GH6 domain; two of these residues (yellow) also line the GH6 binding pocket. **c**, The substrate-binding groove of the GH5 endo-cellulase is shown with residues predicted to interact with cellooctaose (dark blue). The putative catalytic residues — Glu159 (proton donor) and Glu247 (nucleophile) — are highlighted with yellow carbons. Molecular docking suggests an extended cleft accommodating putative +3 and +4 subsites. **d**, The active-site tunnel of the GH6 exo-cellulase is shown with residues predicted to interact with cellohexaose (light blue). Side chains forming the putative catalytic network are highlighted with yellow carbons. Subsites are numbered from the non-reducing to the reducing end as −2, −1, +1, +2, +3, and +4. **e**, Dual-domain GH5–GH6 enzyme exhibits broader substrate specificity than the isolated GH5 catalytic domain (GH5), a CBM-fused GH5 variant (GH5–CBM), or the isolated GH6 catalytic domain (GH6), enabling the hydrolysis of diverse substrates. **f**, HPAEC–PAD chromatograms of soluble products released after incubating 0.5% (w/v) phosphoric acid–swollen cellulose (PASC) with 1 μM enzyme in 50 mM sodium phosphate buffer (pH 7.5) at 30 °C for 12h. **g**, Influence of salinity, temperature, and pH on relative activity of dual-domain GH5–GH6 enzyme. **h**, Synergy experiment combining AA10 lytic polysaccharide monooxygenase (LPMO) with dual-domain GH5–GH6 enzyme show enhanced hydrolytic activity against Avicel. GH5-GH6 each 1 μM were incubated alone or in combination with AA10 for 12 and 24h respectively, all with 1 mM ascorbate included in the reaction mixture.

This evolutionary logic is strikingly mirrored by the host genome: *Lyrodus* ^24^, *Teredo*, and *Xyloredo* (Supplementary Fig. 5) encode multi-domain GH1 cellulases with repeated catalytic modules. Although the precise mechanistic basis for the dual endo- and exo-activities observed in these host-derived enzymes remains to be fully elucidated^24^, both the host and symbiont clearly converge upon a shared evolutionary principle: building larger, more versatile catalytic machines. This structural modularity is a hallmark of biological systems optimized under the stringent ecological constraint of maintaining a low-diversity microbiome.

Biochemical analyses confirm that this structural proximity directly enhances enzymatic efficiency (Fig. 7e,f). Functional assays of heterologously expressed constructs demonstrated that the full-length GH5–GH6 enzyme exhibits substantially higher activity on long-chain cello-oligosaccharides compared to individual domains or CBM-fused variants alone (Fig. 7e,f and Supplementary Fig. 14). Beyond functional synergy, these symbiont cellulases display broad pH and salinity tolerances, consistent with their physical trajectory from gill tissues to the cecum, as well as their constant exposure to seawater ionic strength (Fig. 7g). The enrichment of charged residues within GH5 domains likely confers catalytic halotolerance^55,56^. Together, the enzymatic arsenals of the holobiont exhibit clear signatures of dual optimization for both biochemical efficiency and environmental robustness. Finally, despite stark ecological differences between the shallow-water *T. navalis* and the deep-sea *Xyloredo*, both symbiont genomes retain a nearly identical GH5–GH6–AA10 cellulolytic core (Fig. 7h). This remarkable conservation across >100 million years highlights the strong stabilizing selection that has maintained this fusion-based cellulolytic strategy throughout the entire Teredinidae–Xylophagaidae radiation.

Ultimately, our hologenomic framework redefines the bivalve xylotrophic lifestyle not merely as a marine analog to terrestrial termites, but as a fundamentally distinct evolutionary paradigm for biomass degradation. By trading a hyper-diverse gut microbiome for a highly streamlined, spatially segregated symbiosis, these bivalves achieve greater control over the utilization of the soluble products of lignocellulose degradation than hosts relying on internal microbial fermentation. Instead, through the precise division of metabolic labor, extensive secretory trafficking, and profound structural and enzymatic innovations such as the GH5-GH6 fusion, they have evolved a ’cell-free’ digestive strategy. The extraordinary conservation of this minimalist architecture—across 147 million years and vastly different oceanic depths—highlights its evolutionary robustness, cementing xylotrophic bivalves as uniquely efficient engineers of marine carbon cycling.

## Methods

### Animal sampling

**S**hipworms (*Teredo navalis*) were collected from driftwood in a brackish estuary of the Yellow Sea, China (36°20.40’N, 120°40.80’E). Deep-sea wood-boring clams (*Xyloredo* sp.) were recovered from a depth of ∼1,000 m at the Site F cold seep in the South China Sea (22°06.92’N, 119°17.13’E). *T. navalis* specimens were dissected live. To minimize thermal shock, *Xyloredo* sp. specimens were maintained in chilled seawater during recovery and subsequently dissected onboard. Tissues from both species were flash-frozen for DNA and RNA extraction. All surgical instruments were strictly sterilized between specimens to prevent microbiome cross-contamination. All sampling and experimental procedures were conducted under appropriate permits and in strict compliance with institutional and national guidelines. Voucher materials, including valves, pallets, and tissues, were deposited at the Institute of Marine Science and Technology (IMST), Shandong University, Qingdao, China. Each voucher was assigned a stable accession number, which is reported in Supplementary Table 1.

### FISH and TEM analysis

For fluorescence *in situ* hybridization (FISH) analysis, gill and cecum tissues were fixed in 4% paraformaldehyde in PBS at 4 °C for 16 h, washed three times in cold PBS, dehydrated in 100% methanol, and stored at −20 °C. Tissues were subsequently embedded in Paraplast Plus (Sigma) and cut into 7-μm sections using a Leica microtome. A Cy3-labeled probe targeting the *Teredinibacter* 16S rRNA gene was designed; its specificity was computationally verified against both the host genome and gill metagenomic assemblies to preclude off-target binding. Sequential sections of xylotrophic bivalve tissues were hybridized utilizing the specific probe and its reverse complement (as a negative control), following the protocol described by Halary et al.^57^.

For transmission electron microscopy (TEM), dissected tissues were fixed in 2.5% glutaraldehyde in PBS at 4 °C. Samples were post-fixed in 1% osmium tetroxide and embedded in Epon812 resin. Ultrathin sections (70 nm) generated by a Reichert-Jung ULTRA CUT E ultramicrotome were double-stained with uranyl acetate and lead citrate. The resulting sections were examined using a JEM1400F transmission electron microscope (JEOL) operated at 80 kV.

### Genomic extraction and sequencing

#### DNA extraction

Host genomic DNA was extracted from muscle and mantle tissues, while total DNA for metagenomic analysis was isolated from gill and cecum tissues. All extractions were performed using the DNeasy Blood & Tissue Kit (Qiagen) according to the manufacturer’s protocol. DNA quality and integrity were evaluated via NanoDrop spectrophotometry (Thermo Fisher Scientific) and 1% agarose gel electrophoresis.

#### Genomic sequencing

For host genome assemblies, sequencing libraries were constructed for short-read (Illumina NovaSeq, 150-bp paired-end), long-read (PacBio RSII), and Hi-C (HiSeq X Ten) platforms. Because *Xyloredo* sp. yielded insufficient high-molecular-weight DNA for Hi-C library construction, its genome was assembled only to the contig level. Symbiont metagenomes from gill and cecum tissues were sequenced on the Illumina NovaSeq platform (2 × 150-bp), targeting an average yield of ∼58.22 Gb per sample to ensure deep coverage (Supplementary Tables 2, 3, 7 and 8).

### Genome assembly and annotation

#### Host genome assembly

The host genome was assembled through a combination of PacBio long-read sequencing, Illumina short-read polishing, and Hi-C scaffolding. Initially, PacBio long reads were utilized for *de novo* contig generation via wtdbg2.3^58^. These contigs were then polished with the original PacBio reads using SMRT Link v7.0 (Pacific Biosciences), followed by consensus correction employing Illumina paired-end reads via Pilon v1.22^59^. Redundant haplotypic sequences were subsequently removed with Purge Haplotigs^60^. The polished, haplotig-filtered assembly was then scaffolded to the chromosome scale by integrating Hi-C data with ALLHiC v0.9.13^61^. The final assembly was screened for contamination using FCS-GX v0.5.5 ^62^, and genome completeness was assessed via BUSCO v5.4.4^63^ against the metazoa_odb10 database. Comprehensive sequencing statistics, software versions, and command-line parameters for the host genome assemblies are provided in Supplementary Notes 1 and 2.

#### Repeat annotation

We constructed *de novo* repeat libraries using RepeatModeler 2.0.2 and subsequently identified and soft-masked repeats for both *T. navalis* and *Xyloredo* sp. using RepeatMasker v4.1.7-p1^64^. Sequence divergence from the consensus (Kimura substitution level) and repeat landscapes were generated using the calcDivergenceFromAlign.pl and createRepeatLandscape.pl utilities provided within the RepeatMasker package.

#### Gene prediction and functional annotation

Protein-coding genes were predicted using *ab initio* and transcriptome-based evidence. *Ab initio* gene prediction was performed on the repeat-masked genome using Augustus v3.1^65^, and Genscan^66^ was additionally run using the Silkworm_Genscan.smat parameter file. For transcriptomic evidence, RNA-seq data from various tissues (e.g. mantle, visceral mass, and gill) were mapped to the genome using HISAT2 v2.1.0^67^, and the resulting alignments were assembled into transcripts using StringTie v1.3.4^68^. Candidate open reading frames (ORFs) were predicted utilizing TransDecoder v5.5.0. Transcript-derived ORFs were then projected back to genomic coordinates using cdna_alignment_orf_to_genome_orf.pl. Functional annotations were assigned by querying the UniProt, InterPro, GO, and KEGG databases.

#### Metagenome assembly, binning, and annotation

Gill metagenomic reads were quality-filtered using fastp v0.23.2^69^. Following the *in silico* removal of host-mapped reads, the remaining unmapped reads were assembled using metaSPAdes v3.13.0^70^. The resulting contigs were binned and refined into draft genomes utilizing the ‘binning’, ‘bin_refinement’, and ‘reassemble_bins’ modules of MetaWRAP v1.3.2^71^. Bin completeness and contamination were subsequently assessed using CheckM v1.0.12^72^(Supplementary Tables 7 and 8). To calculate the relative abundance of each bin, clean reads were mapped back to the reassembled bins using SAMtools v1.6^73^. Bacterial gene prediction and functional annotation were performed using Prodigal v2.6.3^74^ and eggNOG-mapper v2.1.12^75,76^, respectively. Finally, the average amino acid identity (AAI) between the recovered bins and symbionts of other xylotrophic bivalves was calculated via the *aai_wf* workflow in CompareM 0.1.2.

#### CAZymes annotation

The *run_dbcan* program within dbCAN3 pipeline^77^ was employed to annotate signature domains for all CAZyme family-including Glycoside Hydrolases (GHs), Glycosyltransferases (GTs), Polysaccharide Lyases (PLs), Carbohydrate Esterases (CEs), Auxiliary Activities (AAs), and Carbohydrate-Binding Modules (CBMs)-across the host and symbiont genomes. Only CAZyme candidates supported by at least two of the three tools (HMMER, DIAMOND, and eCAMI) were retained for further analysis. The assignment of multiple CAZyme domains within a single sequence was permitted. Subsequently, the dbCAN-sub tool was utilized for substrate prediction. Furthermore, all putative lignocelluloses and lignocellulose-binding modules were queried against the dbCAN_sub database using DIAMOND v2.1.8^78^to infer specific enzymatic activities and corroborate their involvement in lignocellulose degradation.

#### RNA extraction and transcriptomic sequencing

Total RNA was extracted from gill and cecum tissues using TRIzol (Invitrogen) and purified with the RNeasy Mini Kit (Qiagen) following the manufacturers’ instructions. RNA integrity and quality were assessed using an Agilent 2100 Bioanalyzer. Metatranscriptomic libraries were constructed from rRNA-depleted RNA (Ribo-Zero Gold) and sequenced on the Illumina NextSeq platform (2×150 bp).

#### Gene expression analysis

Metatranscriptomic reads were mapped to host and symbiont gene models, and expression levels were quantified as transcripts per million (TPM) using Salmon v1.10.3^79^. To normalize host gene expression, mRNA levels were calculated relative to the housekeeping gene *Actin*, a widely utilized internal control in bivalves^80^. Similarly, symbiont gene expression was normalized using *rho*^81^ and *infB*^82^ as internal reference genes. Final symbiont expression values were further standardized based on their relative taxonomic abundance derived from metagenomic data.

#### Phylogenetic analysis and time estimation

To resolve the host phylogenetic position, genomic data from 36 molluscs, two annelids, and one brachiopod species were analyzed (Supplementary Table 19). Single-copy orthologous genes across the assemblies were identified using BUSCO v5.4.4^63^. Amino acid sequences of single-copy BUSCO genes with ≥ 80% taxonomic occupancy were extracted, individually aligned with MAFFT v7.407^83^ under default settings, and trimmed using trimAl v1.4.1^84^ (-gt 0.2). The refined alignments were subsequently concatenated into a phylogenomic supermatrix. A maximum-likelihood (ML) phylogenetic tree was inferred using IQ-TREE v2.2.2.7^85^ under the LG+G4 substitution model, with node support evaluated via 1,000 ultrafast bootstrap replicates. Based on the resulting topology, divergence times were estimated using MCMCTree in PAML v4.9j^86,87^, utilizing fossil records as calibration points (Supplementary Fig. 4). The Markov chain Monte Carlo (MCMC) analysis was run for 10 million generations, with the first 1 million generations discarded as burn-in, and sampled every 1,000 generations to collect 10,000 samples.

For the symbiont phylogeny, all available genomes of the family *Cellvibrionaceae* were retrieved from the NCBI database (Supplementary Table 9). Non-*Cellvibrionaceae* metagenome-assembled genomes (MAGs) recovered in this study were designated as the outgroup. A suite of 120 highly conserved bacterial marker genes was identified, aligned, trimmed, and concatenated using GTDB-Tk v2.3.2^88^ (release 214^89^). The ML phylogeny was constructed using IQ-TREE v2.2.2.7 under default settings, which included automatic substitution model selection, 1,000 ultrafast bootstraps, and 1,000 SH-aLRT replicates^85^.

#### Gene family evolution and functional enrichment

Gene family expansions and contractions were analyzed using CAFE 5^90^ based on orthogroups identified by OrthoFinder v2.5.5^91^. The CAFE analysis was executed in two stages. First, the GAMMA model was utilized to estimate the lambda and alpha parameters, excluding gene families with extreme size variances (>100 copies in a single species). Subsequently, CAFE was run across all families using these fixed parameters to test for evolutionary expansions and contractions. To investigate the genomic basis of shared adaptations in xylotrophic bivalves, we focused on evolutionary events at the most recent common ancestor (MRCA) of *T. navalis* and *Xyloredo* sp. Genes from expanded or contracted families at this focal node were extracted. Gene Ontology (GO) and KEGG pathway enrichment analyses were conducted utilizing the R package clusterProfiler 4.19.7^92^. To account for multiple comparisons, P values were adjusted via the Benjamini–Hochberg false discovery rate (FDR) method, with terms exhibiting an FDR-adjusted P < 0.05 considered significantly enriched.

#### Macrosynteny analysis

Macrosynteny comparisons among the 20 presumed molluscan linkage groups (MLGs), *T. navalis*, and *Xyloredo* sp., along with subsequent data visualization utilizing the R package macrosyntR, were conducted following the methodology described by Sigwart et al.^27^.

#### Homeobox genes identification

Hox and ParaHox genes in representative species of Pteriomorphia and Heterodonta were identified via sequence alignments against the known homeobox gene repertoire of *Mizuhopecten yessoensis* ^93^. Putative gene annotations were subsequently validated through phylogenetic reconstruction.

#### Enzymatic redundancy and signal peptide prediction

To quantitatively assess the redundancy of lignocellulose-degrading enzymes within termite and xylotrophic bivalve holobionts, functional redundancy was evaluated across two classification schemes: Enzyme Commission (EC) numbers and domain-level protein families. For lignocellulases, redundancy metrics were calculated as the ratio of the total number of unique domains to the number of distinct EC numbers or protein families. Additionally, putative secretory signal peptides were predicted using SignalP 6.0^94^.

#### Structural modeling and similarity assessment

The three-dimensional structure of the full-length bi-catalytic GH5-GH6 multi-domain enzyme was predicted utilizing the AlphaFold3 server (https://alphafoldserver.com/). Local Distance Difference Test (LDDT) and TM-score metrics were evaluated for each structural dataset following previous study^95^. Pairwise alignments between structural groups were executed using the Foldseek search module with the parameters -a -e INF --threads 32 --exhaustive-search^96^. Output formats were subsequently customized via the Foldseek convertalis module using the parameters --format-output query,target,evalue,lddt,alntmscore. Mean LDDT and TM-scores were calculated for all pairwise comparisons, strictly excluding intra-group self-alignments.

#### Molecular docking

Putative binding pockets on the enzyme were identified utilizing the DoGSite3 algorithm^97^ via the Proteins*Plus* server (https://proteins.plus)^98^. Cellooctaose and cellohexaose ligands were constructed in ChemDraw v23.1.1. Prior to docking, both receptor and ligand PDB files were converted to the PDBQT format using AutoDockTools v1.5.7^99^. Ligands were docked into the substrate-binding crevice utilizing AutoDock Vina-Carb v1.0^100^. To ensure conformational robustness, docking calculations were performed with an exhaustiveness parameter of 8 and independently repeated 50 times. The top three binding poses based on energetic scores were retained for downstream visualization and analysis in PyMOL.

#### Heterologous expression and enzymatic assays

The substrate specificity and catalytic activities of the purified CAZymes were evaluated against a comprehensive panel of carbohydrates. The tested substrates included crystalline forms (Avicel PH-101 and Whatman filter paper), an amorphous form (phosphoric acid-swollen cellulose, PASC), and various soluble polysaccharides (carboxymethyl cellulose [CMC], β-glucan, xyloglucan, xylan, lichenan, glucomannan, arabinogalactan, and gum arabic). Soluble substrates were dissolved according to the manufacturers’ instructions. Standard reactions were performed in a 300-μL volume containing 50 mM sodium phosphate buffer (pH 7.5) and 0.5 M NaCl. Reaction parameters were tailored based on substrate solubility. For soluble substrates (0.5% w/v), 50 nM of the purified enzyme was added, and the mixture was incubated at 30 °C for 60 min with horizontal agitation (500 rpm). Conversely, assays utilizing insoluble substrates (1% w/v) required 1 μM of enzyme and an extended incubation period of 24 h under identical temperature and agitation conditions. Aliquots were withdrawn at designated intervals. Enzymatic reactions were quenched by the addition of DNS reagent. The release of reducing sugars was subsequently quantified using the standard DNS assay^101^, with glucose serving as the reference standard.

#### Temperature and salinity range of the GH5–GH6 enzyme activity

Standard 300-μL reactions containing 0.5% (w/v) β-glucan and 50 nM of purified enzyme were incubated in a thermomixer for 60 min with horizontal agitation (500 rpm). The tested parameter space encompassed temperatures from 5 to 70 °C, pH levels from 3.0 to 10.0 (maintained utilizing 50 mM sodium citrate for pH 3.0–6.0, potassium phosphate for pH 6.5–8.0, and glycine–NaOH for pH 9.0–10.0), and salinities ranging from 0 to 3.5%. Reactions were immediately quenched with DNS reagent. Product formation was subsequently determined by quantifying the release of reducing sugars against a glucose standard curve, as described (see above).

#### Synergistic cellulose saccharification

The cooperative effect of the GH5–GH6 enzyme and a lytic polysaccharide monooxygenase (LPMO; recombinant AA10 from *Teredinibacter turnerae* T7902) was evaluated. Assays were conducted in 50 mM sodium phosphate buffer (pH 6.0) containing 1% (w/v) crystalline Avicel, supplemented with 1 μM of each enzyme (a 1:1 molar ratio). Enzymatic reactions and subsequent saccharification measurements were performed following the methodology described by Junghare et al. ^102^.

## Data availability

All raw sequencing data (PacBio HiFi, Hi-C, and metagenomic reads) for *T. navalis* and *Xyloredo* sp. have been deposited in the NCBI Sequence Read Archive (SRA) under accession number PRJNA1459029. Assembled host genomes have been deposited in GenBank under project PRJNA1459029. Genome assemblies and annotation files for the hologenomes of both xylotrophic bivalves are freely accessible via Figshare (https://doi.org/10.6084/m9.figshare.32197437). Additionally, the AlphaFold 3-predicted model of the bi-catalytic GH5–GH6 multi-domain enzyme has been deposited in Figshare (https://doi.org/10.6084/m9.figshare.32197437).

## Code availability

Custom code and analytical pipelines utilized in this study have been deposited in Figshare (https://doi.org/10.6084/m9.figshare.32197437).

## Acknowledgements

We thank Dr. J. R. Voight for assistance with species identification, and members of the Y.L. laboratory for their insightful discussions and comments. This work was supported by the National Natural Science Foundation of China (nos. 42376147, 42522609, and 42376088), the Earmarked Fund for CARS (CARS-49), Science & Technology Fundamental Resources Investigation Program (no. 2024FY101000), and the Marine S&T Fund of Shandong Province for Pilot National Laboratory for Marine Science and Technology (Qingdao) (no.2022QNLM030004). Y.L. is supported by the Natural Science Foundation of Shandong Province (ZR2024JQ027). H.S. is supported by the Taishan Scholars Program (no. tsqn202312261), the Excellent Young Scientists Fund of Shandong Province (no. ZR2024YQ005), and the Youth Innovation Promotion Association by CAS (no.2023217). D.L.D is supported by the Gordon and Betty Moore Foundation (https://doi.org/10.37807/GBMF9339) and National Institutes of Health (1R01AI162943-01A1, subaward:10062083-NE).

---

**Extended Data Fig. 1.**
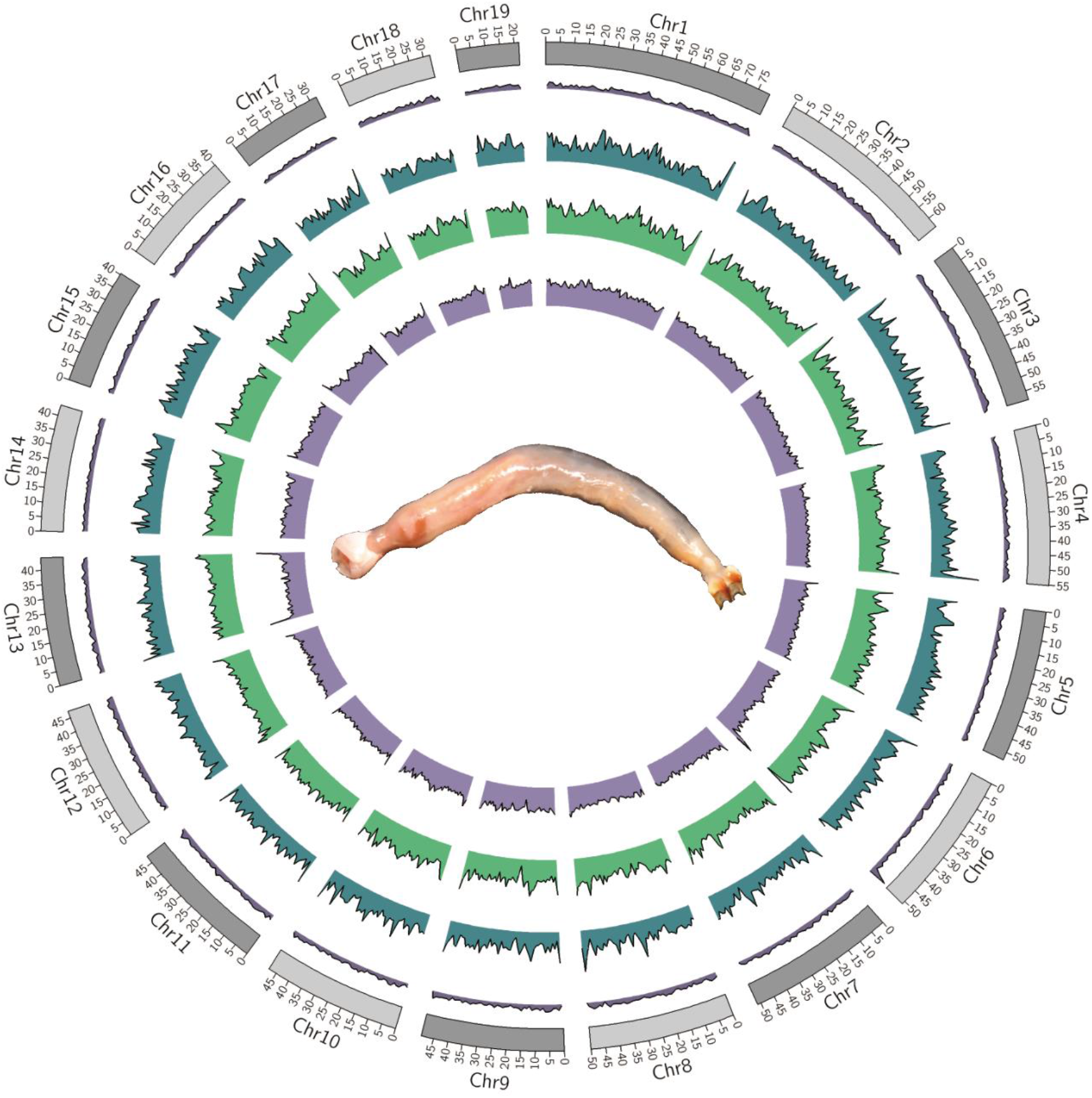
Genome landscape of *Teredo navalis*. From inner to outer rings, the Circos plot displays (1) GC content, (2) repeat sequence density, (3) long terminal repeat (LTR) density, and (4) gene density. All tracks were computed in 1-Mb sliding windows using circos v0.69-8. The center depicts a live *T. navalis* specimen freshly dissected from wood.

**Extended Data Fig. 2.**
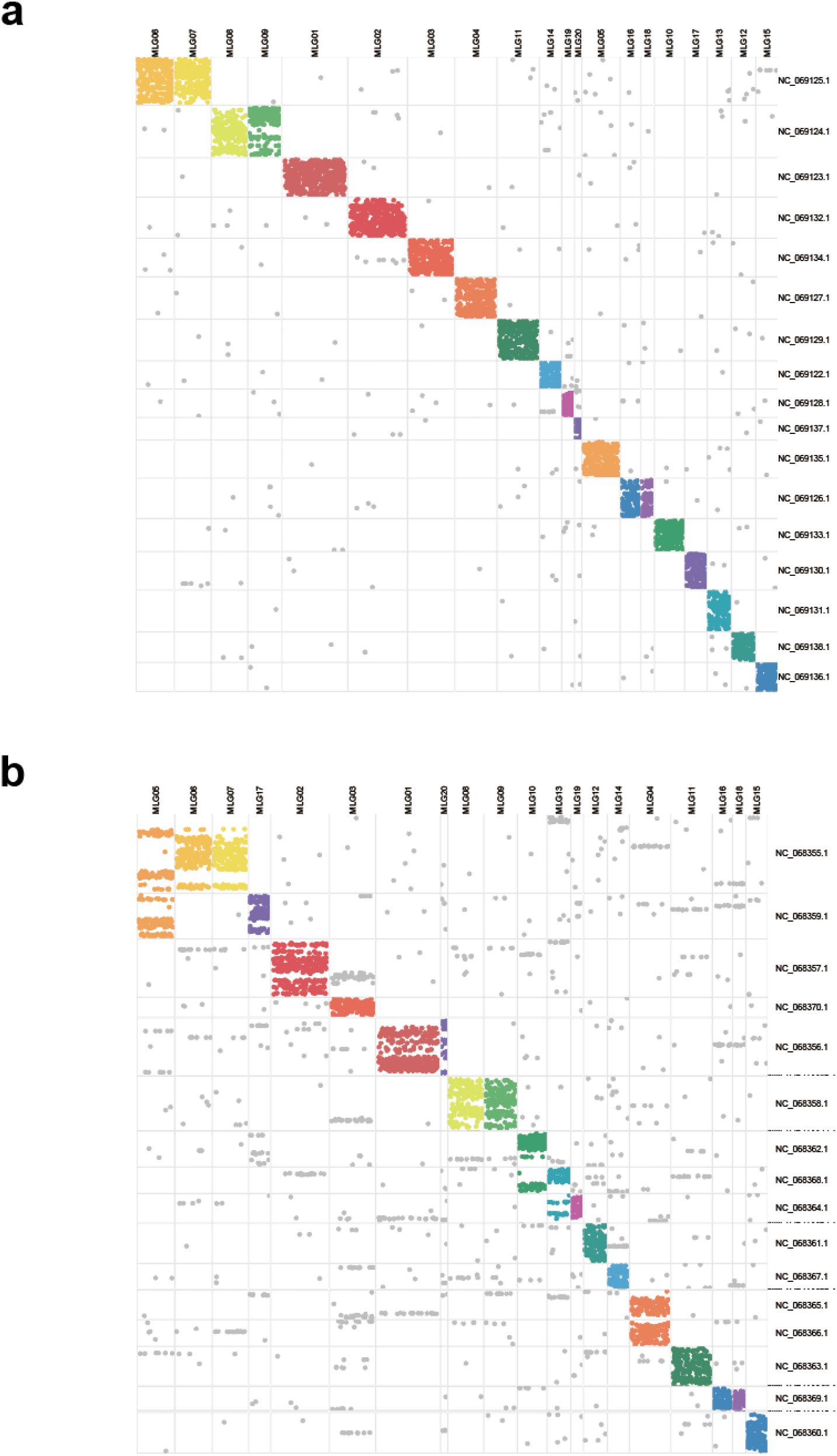
Conserved synteny in close relatives of xylotrophic bivalves. Oxford plots align the genomes of the soft-shell clam (*Mya arenaria*) (**a**) and the zebra mussel (*Dreissena polymorpha*) (**b**) on the y-axis against the ancestral molluscan linkage groups (MLGs) on the x-axis.

**Extended Data Fig. 3.**
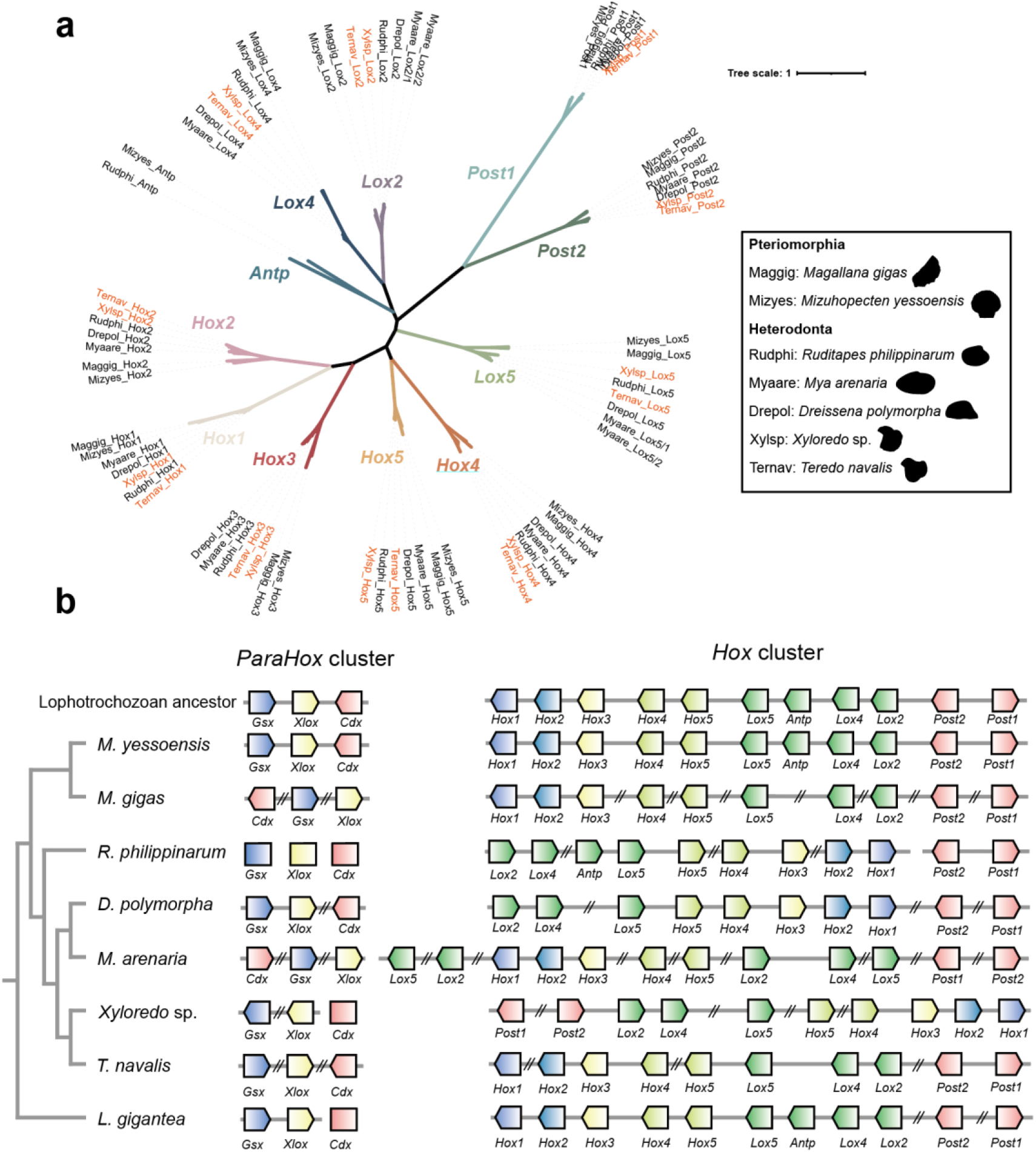
***Hox* and *ParaHox* genes in two xylotrophic bivalves and their close relatives. a**, Unrooted phylogenetic tree of highly conserved Hox proteins across seven bivalve species. Species abbreviations are as follows: Pteriomorphia: *Magallana gigas* (Maggig) and *Mizuhopecten yessoensis* (Mizyes); Heterodonta: *Ruditapes philippinarum* (Rudphi), *Mya arenaria* (Myaare), *Dreissena polymorpha* (Drepol), *Xyloredo* sp. (Xylsp), and *Teredo navalis* (Ternav). The phylogenetic positions of the Hox genes identified in the two xylotrophic bivalves (*T. navalis* and *Xyloredo* sp.) are highlighted in orange. **b**, Schematic of Hox cluster organization in xylotrophic bivalves and other representative bivalves. Solid lines connect genes located on the same scaffold or chromosome (lengths are not proportional to genomic distances). Arrowheads indicate transcriptional orientation, and double slashes denote non-consecutive genes.

**Extended Data Fig. 4.**
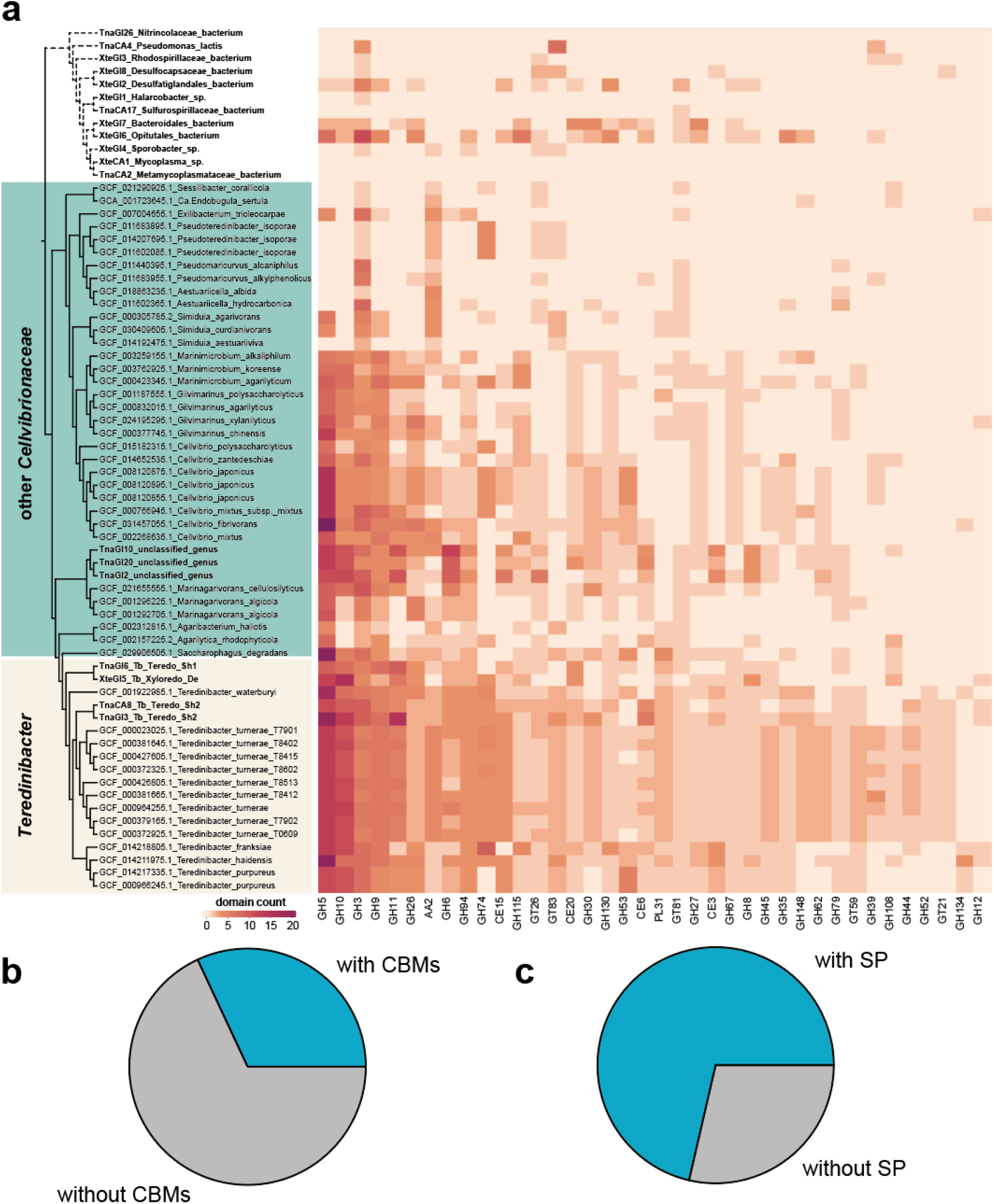
CAZyme family enrichment and phylogenetic relationships within the *Cellvibrionaceae*. **a**, *Teredinibacter* are significantly enriched in specific catalytic CAZyme families compared with other members of the *Cellvibrionaceae*. In the phylogeny, branches corresponding to MAGs recovered in this study are shown in bold, whereas outgroup branches are shown as dashed lines. A substantial proportion of the enriched enzymes in *Teredinibacter* are characterized by the presence of carbohydrate-binding modules (CBMs) and signal peptides (SP), which account for approximately 32% (**b**) and 71% (**c**) of the enzymes, respectively, in the blue section.

**Extended Data Fig. 5.**
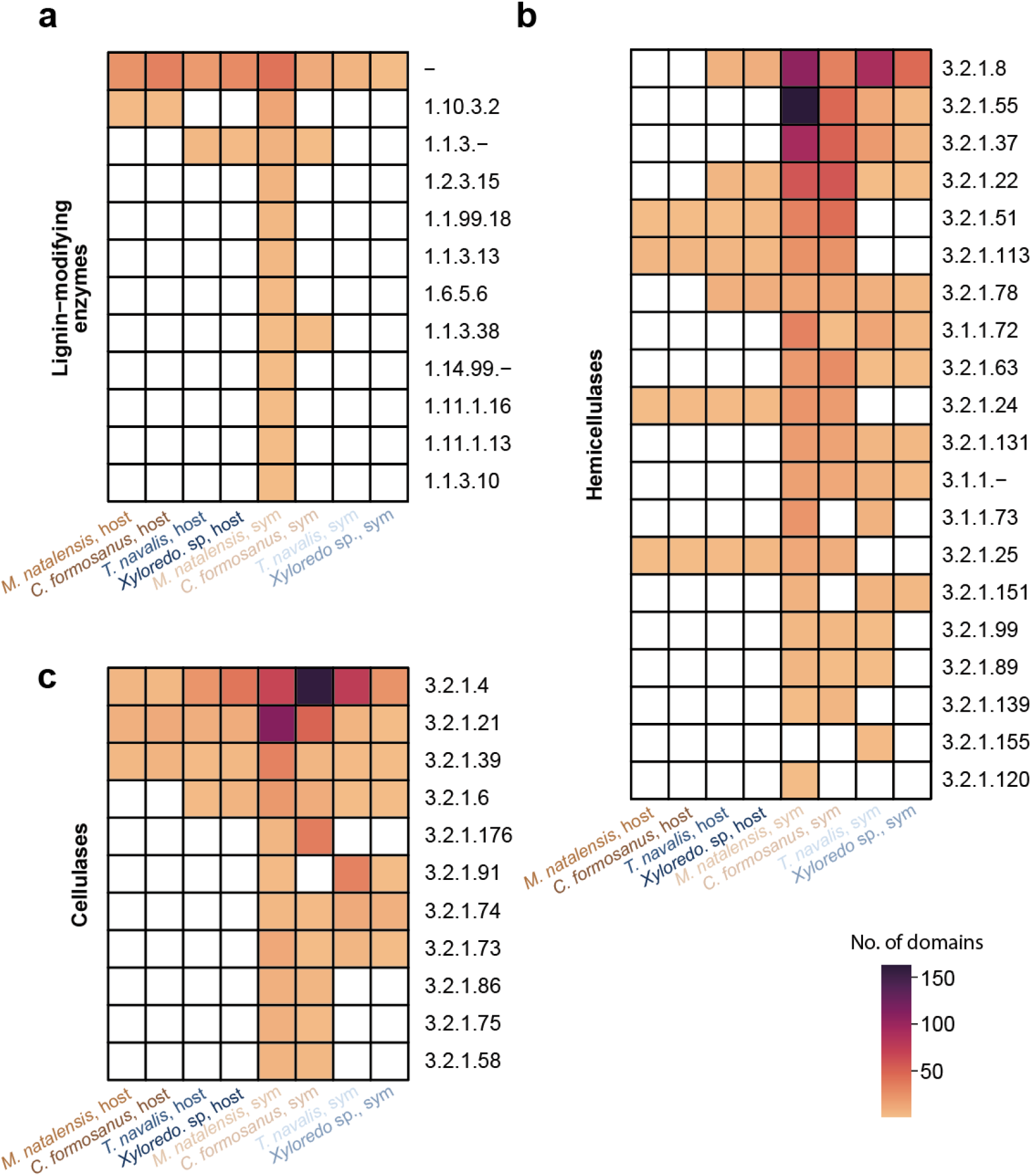
Enzymatic repertoires for wood degradation. EC numbers for lignin-modifying enzymes (**a**), hemicellulases (**b**) and cellulases (**c**) in hosts and symbionts across four symbiotic systems, with the domain count for each EC number.

**Extended Data Fig. 6.**
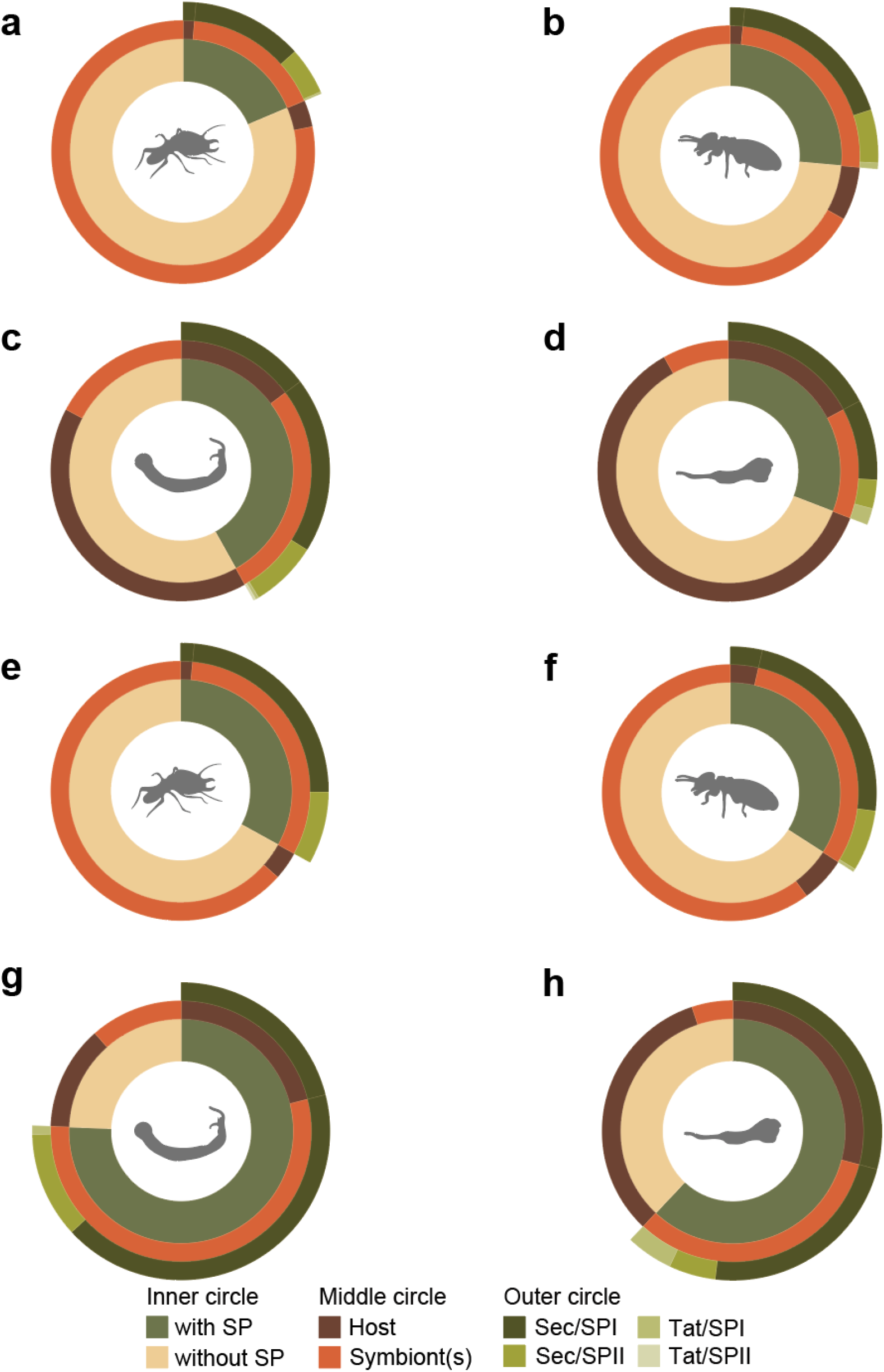
Signal peptide annotation of CAZymes. a–d, Proportion of CAZymes predicted to carry a signal peptide (SP) in each holobiont: **a**, *Macrotermes natalensis*; **b**, *Coptotermes formosanus*; **c**, *Teredo navalis*; **d**, *Xyloredo* sp. e–h, Proportion of lignocellulases predicted to carry a signal peptide in: **e**, *M. natalensis*; **f**, *C. formosanus*; **g**, *T. navalis*; **h**, *Xyloredo* sp.

**Extended Data Fig. 7.**
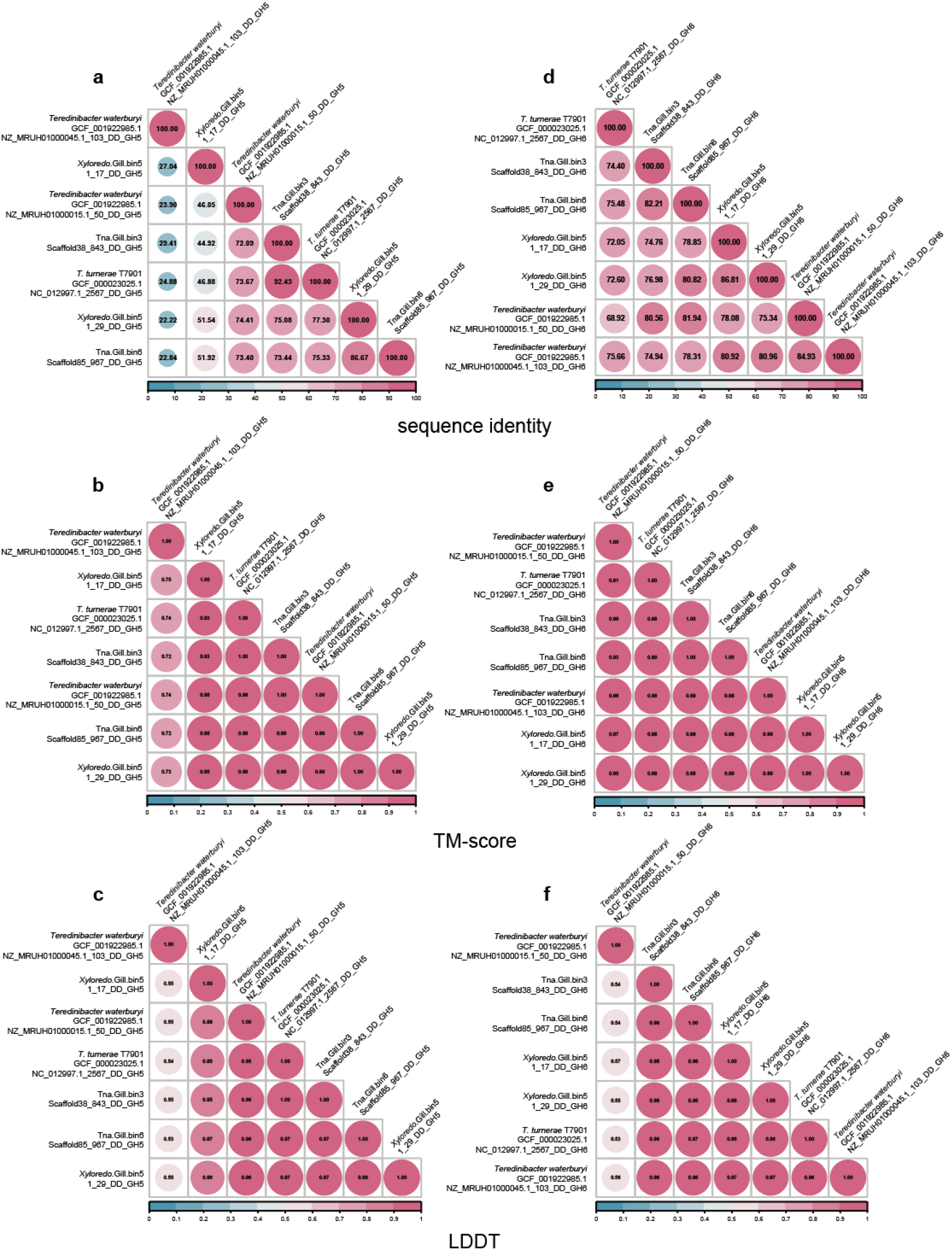
Similarity of GH5 and GH6 catalytic domains within dual-domain GH5–GH6 enzymes across *Teredinibacter* lineages. Sequence identity, TM-score, and LDDT values for the GH5 (**a–c**) and GH6 (**d–f**) domains are presented, respectively.

**Extended Data Fig. 8.**
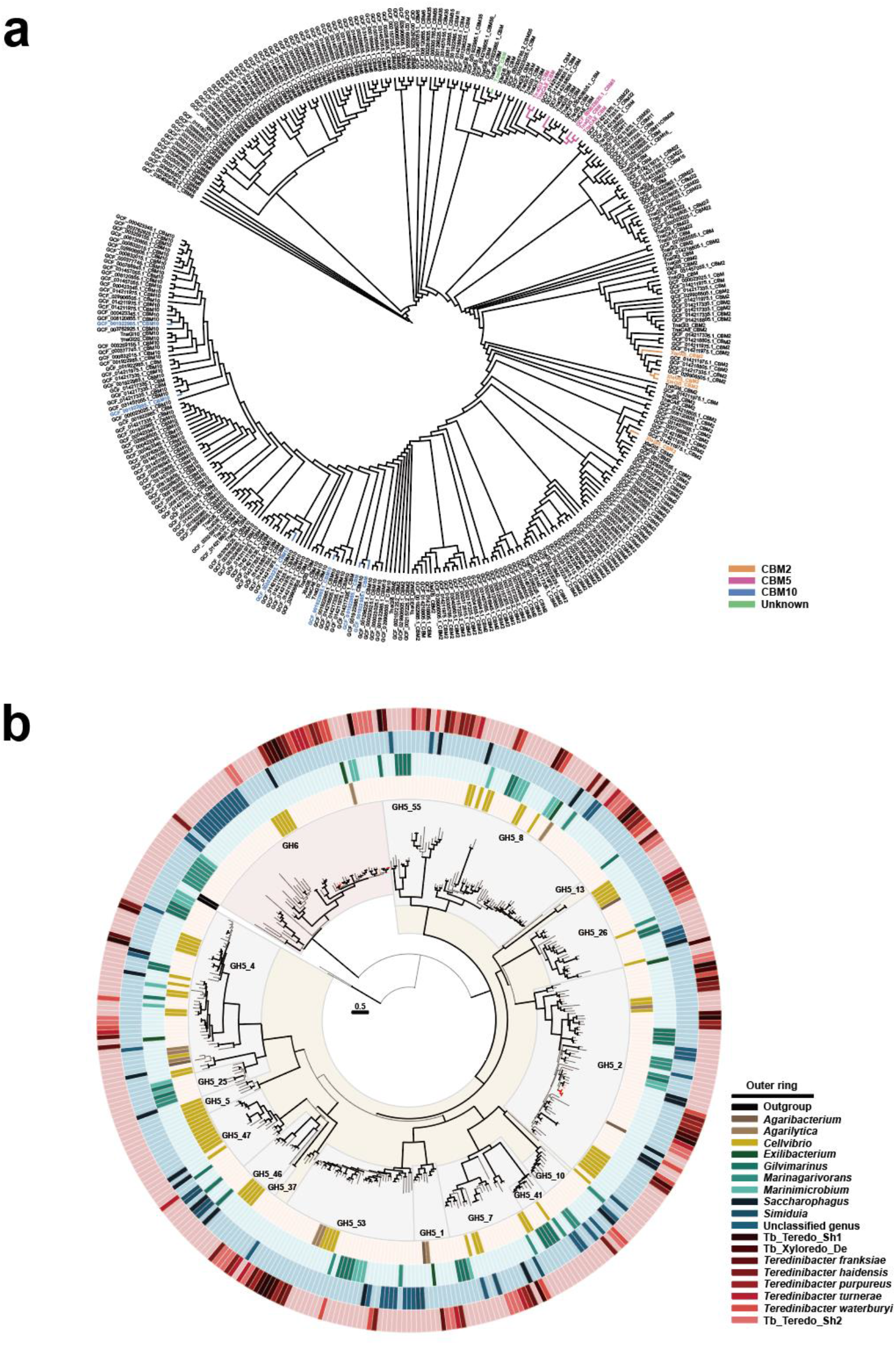
Domain-level phylogenetic analysis of GH5 and GH6 enzymes in *Cellvibrionaceae*. **a**, Phylogenetic tree of 338 GH5 and GH6 domains from *Cellvibrionaceae* (substitution model: Q.pfam+F+R6). Catalytic domains of GH5–GH6 dual-domain enzymes from *Teredinibacter* are highlighted in red. b, Phylogenetic relationships among 337 carbohydrate-binding modules (CBMs) from *Teredinibacter* and closely related taxa. CBMs associated with GH5–GH6 dual-domain enzymes in *Teredinibacter* are highlighted in different colours (substitution model: WAG+F+R4).

## References

1 Bomble, Y. J. et al. Lignocellulose deconstruction in the biosphere. Current opinion in chemical biology 41, 61–70 (2017).

2 Cragg, S. M. et al. Lignocellulose degradation mechanisms across the Tree of Life. Current opinion in chemical biology 29, 108–119 (2015).

3 Johansen, K. S. Lytic polysaccharide monooxygenases: the microbial power tool for lignocellulose degradation. Trends in plant science 21, 926–936 (2016).

4 Hemsworth, G. R., Henrissat, B., Davies, G. J. & Walton, P. H. Discovery and characterization of a new family of lytic polysaccharide monooxygenases. Nature chemical biology 10, 122–126 (2014).

5 Lankiewicz, T. S. et al. Lignin deconstruction by anaerobic fungi. Nature Microbiology 8, 596–610 (2023).

6 Hsin, K.-T. et al. Lignocellulose degradation in bacteria and fungi: cellulosomes and industrial relevance. Frontiers in Microbiology 16, 1583746 (2025).

7 Varma, A., Kolli, B. K., Paul, J., Saxena, S. & König, H. Lignocellulose degradation by microorganisms from termite hills and termite guts: a survey on the present state of art. FEMS Microbiology Reviews 15, 9–28 (1994).

8 Voight, J. R. Xylotrophic bivalves: aspects of their biology and the impacts of humans. Journal of Molluscan Studies 81, 175–186 (2015).

9 Pournou, A. in Biodeterioration of Wooden Cultural Heritage: Organisms and Decay Mechanisms in Aquatic and Terrestrial Ecosystems 261–343 (Springer, 2020).

10 Popham, J. & Dickson, M. Bacterial associations in the teredo Bankia australis (Lamellibranchia: Mollusca). Marine Biology 19, 338–340 (1973).

11 Popham, J. Further observations of the gland of Deshayes in the teredo Bankia australis. Veliger 18, 55–59 (1974).

12 Distel, D. L., Beaudoin, D. J. & Morrill, W. Coexistence of multiple proteobacterial endosymbionts in the gills of the wood-boring bivalve Lyrodus pedicellatus (Bivalvia: Teredinidae). Applied and Environmental Microbiology 68, 6292–6299 (2002).

13 Yang, J. C. et al. The complete genome of Teredinibacter turnerae T7901: an intracellular endosymbiont of marine wood-boring bivalves (shipworms). PloS one 4, e6085 (2009).

14 O’Connor, R. M. et al. Gill bacteria enable a novel digestive strategy in a wood-feeding mollusk. Proceedings of the National Academy of Sciences 111, E5096–E5104 (2014).

15 Distel, D. L., Morrill, W., MacLaren-Toussaint, N., Franks, D. & Waterbury, J. Teredinibacter turnerae gen. nov., sp. nov., a dinitrogen-fixing, cellulolytic, endosymbiotic gamma-proteobacterium isolated from the gills of wood-boring molluscs (Bivalvia: Teredinidae). International Journal of Systematic and Evolutionary Microbiology 52, 2261–2269 (2002).

16 Altamia, M. A. & Distel, D. L. Transport of symbiont-encoded cellulases from the gill to the gut of shipworms via the enigmatic ducts of Deshayes: a 174- year mystery solved. Proceedings of the Royal Society B 289, 20221478 (2022).

17 Distel, D. L. & Roberts, S. J. Bacterial endosymbionts in the gills of the deep-sea wood-boring bivalves Xylophaga atlantica and Xylophaga washingtona. The Biological Bulletin 192, 253–261 (1997).

18 Turner, R. D. On the subfamily Xylophagainae (Family Pholadidae, Bivalvia, Mollusca). Bulletin of the Museum of Comparative Zoology 157, 223–308 (2002).

19 Distel, D. L. et al. Molecular phylogeny of Pholadoidea Lamarck, 1809 supports a single origin for xylotrophy (wood feeding) and xylotrophic bacterial endosymbiosis in Bivalvia. Molecular phylogenetics and evolution 61, 245–254 (2011).

20 Romano, C. et al. Wooden stepping stones: diversity and biogeography of deep-sea wood boring Xylophagaidae (Mollusca: Bivalvia) in the North-East Atlantic Ocean, with the description of a new genus. Frontiers in Marine Science 7, 579959 (2020).

21 Fagervold, S. K. et al. Microbial communities in sunken wood are structured by wood-boring bivalves and location in a submarine canyon. PloS one 9, e96248 (2014).

22 Wardman, J. F., Bains, R. K., Rahfeld, P. & Withers, S. G. Carbohydrate-active enzymes (CAZymes) in the gut microbiome. Nature Reviews Microbiology 20, 542–556 (2022).

23 Mootz, H. D., Schwarzer, D. & Marahiel, M. A. Construction of hybrid peptide synthetases by module and domain fusions. Proceedings of the National Academy of Sciences 97, 5848–5853 (2000).

24 Sabbadin, F. et al. Uncovering the molecular mechanisms of lignocellulose digestion in shipworms. Biotechnology for biofuels 11, 59 (2018).

25 Betcher, M. A. et al. Microbial distribution and abundance in the digestive system of five shipworm species (Bivalvia: Teredinidae). (2012).

26 Distel, D. L. The biology of marine wood boring bivalves and their bacterial endosymbionts. (2003).

27 Sigwart, J. D., Li, Y., Chen, Z., Vonc ina, K. & Sun, J. Still waters run deep in large-scale genome rearrangements of morphologically conservative Polyplacophora. Elife 13, RP102542 (2025).

28 Zhang, Y. et al. Comparative genomics reveals evolutionary drivers of sessile life and left-right shell asymmetry in bivalves. Genomics, proteomics & bioinformatics 20, 1078–1091 (2022).

29 Gruffydd, L. D., Lane, D. & Beaumont, A. The glands of the larval foot in Pecten maximus L. and possible homologues in other bivalves. Journal of the Marine Biological Association of the United Kingdom 55, 463–476 (1975).

30 Robin, N. et al. The oldest shipworms (Bivalvia, Pholadoidea, Teredinidae) preserved with soft parts (western France): insights into the fossil record and evolution of Pholadoidea. Palaeontology 61, 905–918 (2018).

31 Gasser, M. T. et al. Membrane vesicles can contribute to cellulose degradation by Teredinibacter turnerae, a cultivable intracellular endosymbiont of shipworms. Microbial Biotechnology 17, e70064 (2024).

32 McCutcheon, J. P. & Moran, N. A. Extreme genome reduction in symbiotic bacteria. Nature Reviews Microbiology 10, 13–26 (2012).

33 Wernegreen, J. J. Genome evolution in bacterial endosymbionts of insects. Nature Reviews Genetics 3, 850–861 (2002).

34 Luyten, Y. A., Thompson, J. R., Morrill, W., Polz, M. F. & Distel, D. L. Extensive variation in intracellular symbiont community composition among members of a single population of the wood-boring bivalve Lyrodus pedicellatus (Bivalvia: Teredinidae). Applied and environmental microbiology 72, 412–417 (2006).

35 Speare, L. & Distel, D. L. Cultivation and Fluorescent in situ hybridization suggest that some shipworm species acquire endosymbiotic bacteria through indirect horizontal transmission. bioRxiv, 2022.2011. 2013.516348 (2022).

36 Stravoravdis, S., Shipway, J. R. & Goodell, B. How do shipworms eat wood? Screening shipworm gill symbiont genomes for lignin-modifying enzymes. Frontiers in Microbiology 12, 665001 (2021).

37 Janusz, G. et al. Lignin degradation: microorganisms, enzymes involved, genomes analysis and evolution. FEMS microbiology reviews 41, 941–962 (2017).

38 Bredon, M., Dittmer, J., Noël, C., Moumen, B. & Bouchon, D. Lignocellulose degradation at the holobiont level: teamwork in a keystone soil invertebrate. Microbiome 6, 162 (2018).

39 Brune, A. Symbiotic digestion of lignocellulose in termite guts. Nature Reviews Microbiology 12, 168–180 (2014).

40 Wang, K. et al. Lignocellulose degradation in Protaetia brevitarsis larvae digestive tract: refining on a tightly designed microbial fermentation production line. Microbiome 10, 90 (2022).

41 Ohkuma, M. Termite symbiotic systems: efficient bio-recycling of lignocellulose. Applied microbiology and biotechnology 61, 1–9 (2003).

42 Auer, L. et al. Uncovering the potential of termite gut microbiome for lignocellulose bioconversion in anaerobic batch bioreactors. Frontiers in microbiology 8, 2623 (2017).

43 Liu, N. et al. Functional metagenomics reveals abundant polysaccharide-degrading gene clusters and cellobiose utilization pathways within gut microbiota of a wood-feeding higher termite. The ISME journal 13, 104–117 (2019).

44 Lo, N., Tokuda, G. & Watanabe, H. in Biology of Termites: a modern synthesis 51–67 (Springer, 2010).

45 Cann, I. et al. Rumen-targeted mining of enzymes for bioenergy production. Annual Review of Animal Biosciences 13 (2025).

46 Waterbury, J. B., Calloway, C. B. & Turner, R. D. A cellulolytic nitrogen-fixing bacterium cultured from the gland of Deshayes in shipworms (Bivalvia: Teredinidae). Science 221, 1401–1403 (1983).

47 Turner, R. D. Wood-boring bivalves, opportunistic species in the deep sea. Science 180, 1377–1379 (1973).

48 Freer, S., Greene, R. & Bothast, R. (ACS Publications, 2001).

49 Ekborg, N. A., Morrill, W., Burgoyne, A. M., Li, L. & Distel, D. L. CelAB, a multifunctional cellulase encoded by Teredinibacter turnerae T7902T, a culturable symbiont isolated from the wood-boring marine bivalve Lyrodus pedicellatus. Applied and environmental microbiology 73, 7785–7788 (2007).

50 Howard, M. B., Ekborg, N. A., Taylor, L. E., Hutcheson, S. W. & Weiner, R. M. Identification and analysis of polyserine linker domains in prokaryotic proteins with emphasis on the marine bacterium Microbulbifer degradans. Protein science 13, 1422–1425 (2004).

51 Shen, H. et al. Deletion of the linker connecting the catalytic and cellulose- binding domains of endoglucanase A (CenA) of Cellulomonas fimi alters its conformation and catalytic activity. Journal of Biological Chemistry 266, 11335–11340 (1991).

52 El-Gendi, H. et al. A comprehensive insight into fungal enzymes: structure, classification, and their role in mankind’s challenges. Journal of Fungi 8, 23 (2021).

53 Sero, E. T. et al. Exploring the termite gut as a hub of industrially important microbes and enzymes for biofuel production. Biomass and Bioenergy 199, 107899 (2025).

54 Zhou, M. et al. Recent progress of dual-site catalysts in emerging electrocatalysis: a review. Catalysis Science & Technology 13, 4615–4634 (2023).

55 Ghosh, D. et al. A Highly Thermostable and Novel GH5 Endoglucanase from Bacillus sp. Strain BS with Enhanced Biomass Saccharification Potential in Seawater. bioRxiv, 2025.2012. 2021.695781 (2025).

56 Zarafeta, D. et al. Discovery and characterization of a thermostable and highly halotolerant GH5 cellulase from an icelandic hot spring isolate. PLoS One 11, e0146454 (2016).

57 Halary, S., Riou, V., Gaill, F., Boudier, T. & Duperron, S. 3D FISH for the quantification of methane-and sulphur-oxidizing endosymbionts in bacteriocytes of the hydrothermal vent mussel Bathymodiolus azoricus. The ISME Journal 2, 284–292 (2008).

58 Ruan, J. & Li, H. Fast and accurate long-read assembly with wtdbg2. Nature methods 17, 155–158 (2020).

59 Walker, B. J. et al. Pilon: an integrated tool for comprehensive microbial variant detection and genome assembly improvement. PloS one 9, e112963 (2014).

60 Roach, M. J., Schmidt, S. A. & Borneman, A. R. Purge Haplotigs: allelic contig reassignment for third-gen diploid genome assemblies. BMC bioinformatics 19, 460 (2018).

61 Zhang, X., Zhang, S., Zhao, Q., Ming, R. & Tang, H. Assembly of allele-aware, chromosomal-scale autopolyploid genomes based on Hi-C data. Nature plants 5, 833–845 (2019).

62 Astashyn, A. et al. Rapid and sensitive detection of genome contamination at scale with FCS-GX. Genome biology 25, 60 (2024).

63 Manni, M., Berkeley, M. R., Seppey, M., Simão, F. A. & Zdobnov, E. M. BUSCO update: novel and streamlined workflows along with broader and deeper phylogenetic coverage for scoring of eukaryotic, prokaryotic, and viral genomes. Molecular biology and evolution 38, 4647–4654 (2021).

64 Flynn, J. M. et al. RepeatModeler2 for automated genomic discovery of transposable element families. Proceedings of the National Academy of Sciences 117, 9451–9457 (2020).

65 Stanke, M. et al. AUGUSTUS: ab initio prediction of alternative transcripts. Nucleic acids research 34, W435–W439 (2006).

66 Burge, C. & Karlin, S. Prediction of complete gene structures in human genomic DNA. Journal of molecular biology 268, 78–94 (1997).

67 Kim, D., Paggi, J. M., Park, C., Bennett, C. & Salzberg, S. L. Graph-based genome alignment and genotyping with HISAT2 and HISAT-genotype. Nature biotechnology 37, 907–915 (2019).

68 Pertea, M. et al. StringTie enables improved reconstruction of a transcriptome from RNA-seq reads. Nature biotechnology 33, 290–295 (2015).

69 Chen, S., Zhou, Y., Chen, Y. & Gu, J. fastp: an ultra-fast all-in-one FASTQ preprocessor. Bioinformatics 34, i884–i890 (2018).

70 Nurk, S., Meleshko, D., Korobeynikov, A. & Pevzner, P. A. metaSPAdes: a new versatile metagenomic assembler. Genome research 27, 824–834 (2017).

71 Uritskiy, G. V., DiRuggiero, J. & Taylor, J. MetaWRAP—a flexible pipeline for genome-resolved metagenomic data analysis. Microbiome 6, 158 (2018).

72 Parks, D. H., Imelfort, M., Skennerton, C. T., Hugenholtz, P. & Tyson, G. W. CheckM: assessing the quality of microbial genomes recovered from isolates, single cells, and metagenomes. Genome research 25, 1043–1055 (2015).

73 Danecek, P. et al. Twelve years of SAMtools and BCFtools. Gigascience 10, giab008 (2021).

74 Hyatt, D. et al. Prodigal: prokaryotic gene recognition and translation initiation site identification. BMC bioinformatics 11, 1–11 (2010).

75 Cantalapiedra, C. P., Hernández-Plaza, A., Letunic, I., Bork, P. & Huerta-Cepas, J. eggNOG-mapper v2: functional annotation, orthology assignments, and domain prediction at the metagenomic scale. Molecular biology and evolution 38, 5825–5829 (2021).

76 Huerta-Cepas, J. et al. eggNOG 5.0: a hierarchical, functionally and phylogenetically annotated orthology resource based on 5090 organisms and 2502 viruses. Nucleic acids research 47, D309–D314 (2019).

77 Zheng, J. et al. dbCAN3: automated carbohydrate-active enzyme and substrate annotation. Nucleic acids research 51, W115–W121 (2023).

78 Buchfink, B., Reuter, K. & Drost, H.-G. Sensitive protein alignments at tree-of-life scale using DIAMOND. Nature methods 18, 366–368 (2021).

79 Patro, R., Duggal, G., Love, M. I., Irizarry, R. A. & Kingsford, C. Salmon provides fast and bias-aware quantification of transcript expression. Nature methods 14, 417–419 (2017).

80 Iuffrida, L. et al. Stability and expression patterns of housekeeping genes in Mediterranean mussels (Mytilus galloprovincialis) under field investigations. Comparative Biochemistry and Physiology Part C: Toxicology & Pharmacology 287, 110047 (2025).

81 Gomes, A. É. I. et al. Selection and validation of reference genes for gene expression studies in Klebsiella pneumoniae using Reverse Transcription Quantitative real-time PCR. Scientific reports 8, 9001 (2018).

82 Laalami, S., Putzer, H., Plumbridge, J. A. & Grunberg-Manago, M. A severely truncated form of translational initiation factor 2 supports growth of Escherichia coli. Journal of molecular biology 220, 335–349 (1991).

83 Katoh, K. & Standley, D. M. MAFFT multiple sequence alignment software version 7: improvements in performance and usability. Molecular biology and evolution 30, 772–780 (2013).

84 Capella-Gutiérrez, S., Silla-Martínez, J. M. & Gabaldón, T. trimAl: a tool for automated alignment trimming in large-scale phylogenetic analyses. Bioinformatics 25, 1972–1973 (2009).

85 Minh, B. Q. et al. IQ-TREE 2: new models and efficient methods for phylogenetic inference in the genomic era. Molecular biology and evolution 37, 1530–1534 (2020).

86 Yang, Z. PAML: a program package for phylogenetic analysis by maximum likelihood. Bioinformatics 13, 555–556 (1997). 10.1093/bioinformatics/13.5.555

87 Yang, Z. PAML 4: Phylogenetic Analysis by Maximum Likelihood. Molecular Biology and Evolution 24, 1586–1591 (2007). 10.1093/molbev/msm088

88 Chaumeil, P.-A., Mussig, A. J., Hugenholtz, P. & Parks, D. H. GTDB-Tk v2: memory friendly classification with the genome taxonomy database. Bioinformatics 38, 5315–5316 (2022). 10.1093/bioinformatics/btac672

89 Parks, D. H. et al. GTDB: an ongoing census of bacterial and archaeal diversity through a phylogenetically consistent, rank normalized and complete genome-based taxonomy. Nucleic Acids Research 50, D785–D794 (2021). 10.1093/nar/gkab776

90 Mendes, F. K., Vanderpool, D., Fulton, B. & Hahn, M. W. CAFE 5 models variation in evolutionary rates among gene families. Bioinformatics 36, 5516–5518 (2020). 10.1093/bioinformatics/btaa1022

91 Emms, D. M. & Kelly, S. OrthoFinder: phylogenetic orthology inference for comparative genomics. Genome biology 20, 238 (2019).

92 Wu, T. et al. clusterProfiler 4.0: A universal enrichment tool for interpreting omics data. The innovation 2 (2021).

93 Wang, S. et al. Scallop genome provides insights into evolution of bilaterian karyotype and development. Nature ecology & evolution 1, 0120 (2017).

94 Teufel, F. et al. SignalP 6.0 predicts all five types of signal peptides using protein language models. Nature biotechnology 40, 1023–1025 (2022).

95 Wu, W. et al. Structural genomics sheds light on protein functions and remote homologs across the insect tree of life. Cell Research, 1–15 (2026).

96 Van Kempen, M. et al. Fast and accurate protein structure search with Foldseek. Nature biotechnology 42, 243–246 (2024).

97 Graef, J., Ehrt, C. & Rarey, M. Binding site detection remastered: enabling fast, robust, and reliable binding site detection and descriptor calculation with DoGSite3. Journal of Chemical Information and Modeling 63, 3128–3137 (2023).

98 Ehrt, C. et al. ProteinsPlus: a publicly available resource for protein structure mining. Nucleic Acids Research 53, W478–W484 (2025). 10.1093/nar/gkaf377

99 Morris, G. M. et al. AutoDock4 and AutoDockTools4: Automated docking with selective receptor flexibility. Journal of computational chemistry 30, 2785–2791 (2009).

100 Nivedha, A. K., Thieker, D. F., Makeneni, S., Hu, H. & Woods, R. J. Vina-Carb: improving glycosidic angles during carbohydrate docking. Journal of chemical theory and computation 12, 892–901 (2016).

101 Miller, G. L. Use of dinitrosalicylic acid reagent for determination of reducing sugar. Analytical chemistry 31, 426–428 (1959).

102 Junghare, M. et al. Biochemical and structural characterisation of a family GH5 cellulase from endosymbiont of shipworm P. megotara. Biotechnology for Biofuels and Bioproducts 16, 61 (2023).

